# Structure and Function of the Dot/Icm T4SS

**DOI:** 10.1101/2023.03.22.533729

**Authors:** Przemysław Dutka, Yuxi Liu, Stefano Maggi, Debnath Ghosal, Jue Wang, Stephen D. Carter, Wei Zhao, Sukhithasri Vijayrajratnam, Joseph P. Vogel, Grant J. Jensen

## Abstract

The *Legionella pneumophila* Dot/Icm type IV secretion system (T4SS) delivers effector proteins into host cells during infection. Despite its significance as a potential drug target, our current understanding of its atomic structure is limited to isolated subcomplexes. In this study, we used subtomogram averaging and integrative modeling to construct a nearly-complete model of the Dot/Icm T4SS accounting for seventeen protein components. We locate and provide insights into the structure and function of six new components including DotI, DotJ, DotU, IcmF, IcmT, and IcmX. We find that the cytosolic N-terminal domain of IcmF, a key protein forming a central hollow cylinder, interacts with DotU, providing insight into previously uncharacterized density. Furthermore, our model, in combination with analyses of compositional heterogeneity, explains how the cytoplasmic ATPase DotO is connected to the periplasmic complex via interactions with membrane-bound DotI/DotJ proteins. Coupled with *in situ* infection data, our model offers new insights into the T4SS-mediated secretion mechanism.

## Introduction

Bacterial type IV secretion systems (T4SS) are among the largest macromolecular complexes known and are present in most classes of bacteria (*1*). T4SSs are used to translocate DNA and protein substrates across bacterial cell envelopes, often into other target cells in a contact-dependent manner (*2*). For example, conjugative T4SSs drive bacterial conjugation by facilitating DNA transfer, an important process in the spread of antibiotic resistance (*3*).

*Legionella pneumophila* is a Gram-negative bacterial pathogen that causes a form of pneumonia called Legionnaires’ disease. There have been 10,000 reported cases of Legionnaires’ disease in the USA so far with a 10% fatality rate (*4*). *L. pneumophila* causes disease by surviving and replicating within alveolar macrophages (*5, 6*). Intracellular survival is mediated by a specialized T4SS known as Dot/Icm (defective in organelle trafficking/intracellular multiplication). The Dot/Icm system exports over 300 effector proteins into host cells that modulate the host’s cell biology, including its histone modification, transcriptome, proteome, and vesicle trafficking (*7*). This results in the prevention of phagosome-lysosome fusion and the establishment of a vacuole permissive for bacterial replication.

*In situ* (*8–13*) and *in vitro* (*14–17*) structural studies of T4SSs have revealed their complexity and elegance. Our early cryo-electron tomography (cryo-ET) studies (*8, 10*), and those of others (*12, 13*), revealed the overall architecture of the intact Dot/Icm T4SS *in situ* and began to dissect the molecular organization of this complex (*10, 12*). Single particle reconstructions then resolved the outer membrane complex (OMC) and periplasmic ring (PR) to near-atomic resolution, identifying new components and symmetry mismatches between the OMC and PR (*15, 16*).

To provide a more comprehensive understanding of the system, here we collected an order of magnitude more data and performed extensive subtomogram averaging with a focus on mobile connections within the system. Exploiting the improved quality of our maps, new structural and biochemical data, and advances in protein structure predictions, we construct a nearly complete molecular model of the *L. pneumophila* Dot/Icm T4SS. Our model incorporates over 460,000 atoms, more than double the number of atoms in any previous model of the system (*10, 12, 13, 16*). Together with an analysis of compositional heterogeneity and infection *in situ*, our model sheds light on the T4SS-mediated secretion mechanism. We particularly highlight the role of IcmF as a primary component that forms a central hollow cylinder. We show that IcmF is a flexible protein spanning the inner membrane (IM) with a Ras-like domain on the cytosolic site. IcmF, and its interacting partner DotU, play a crucial role in polar targeting (*10*), biogenesis (*10*) and secretion of effectors (*18*). Furthermore, we show that the cytosolic ATPases connect with the periplasmic complex via a linker domain formed by DotI:DotJ complexes and the collar density, connections we propose form an important signaling pathway between the cytoplasm and the OMC.

## Results

### Compositional Heterogeneity of the Dot/Icm T4SS

To further characterize the T4SS beyond our previous work (*10*), we collected a much larger dataset of approximately 900 tomograms, resulting in ~5,700 particles (Fig. S1). Because our previous work had shown that a fusion of superfolder green fluorescent protein (sfGFP) to the C-terminus of DotF (DotF-sfGFP) stabilizes the complex (*10*), we again imaged this strain. We also imaged a derivative strain with an additional DotB (E191K) mutation, which locks the cytoplasmic ATPase DotB onto membrane-associated DotO ATPase (*19*)(*8, 10, 12*), (*19*). Subtomogram averages were produced using data combined from both strains.

One notable feature of the Dot/Icm T4SS and related systems is the unique architecture of the cytoplasmic ATPases (*12, 17*). Typically, AAA+ ATPases assemble into hexameric structures, but in T4SS, DotO and its homologs (e.g. VirB4) form a unique hexamer of dimers. Each copy of DotO in a dimer is in a different conformation (*12, 17*). In our previous work, the structure of the cytoplasmic complex was unclear, possibly due to compositional heterogeneity (*8*, *10*). Here we conducted extensive 3D classification to gain a better understanding of the composition and heterogeneity of the inner membrane complex (IMC) *in situ* (Fig. S2). After testing several programs, we obtained the best results using the I3 subtomogram averaging package (*20*).

We performed 3D classification using a simple cylindrical mask enclosing the entire T4SS. This approach allowed us to classify a fraction of the particles that lack the IMC (14%, Class I) (Fig. S2A). We then used the remaining particles to perform focused refinement and classification on the IMC. This approach revealed two additional classes of particles. In the largest class (72%, Class II), the arrangement of the ATPases was ambiguous. Although we could resolve a central barrel surrounded by external densities, their connectivity and symmetry was unclear. This class might reflect either a partially assembled state or misalignment of particles. The final class, which contained only 14% (Class III) of the original data set, exhibited the previously described hexamer-of-dimers arrangement (*12*). Notably, the presence of the hexamer of dimers was associated with a change in the IM curvature. However, no difference in the distance between the IM and outer membrane (OM) was observed, regardless of the cytoplasmic complex composition.

To confirm our classification results we analyzed individual particles imaged in thin *L. pneumophila* cell lamella prepared by cryogenic focused-ion-beam milling (cryo-FIB-milling). Substantial heterogeneity in the cytoplasmic complex was observed (Fig. S2B). We identified particles without IMC, with asymmetric IMC density, and with characteristic V-shaped IMC density connecting to the collar via the linker domain, confirming our 3D classes.

After performing focused refinement on the IMC in particle classes II and III, 13-fold symmetry was no longer apparent in the OMC, indicating that the periplasmic and cytoplasmic densities are not rotationally linked. To analyze the potential stabilizing effect of the linker connecting with the collar, we performed another round of focused refinement with a mask around the periplasmic complex and examined how much the particles moved. We found that particles shifted substantially more in class II where the linker and outer ATPases were not well-resolved (Fig. S2C) compared to class III where the linker, outer ATPases and collar were better resolved (Fig. S2D). In both cases clear s-fold symmetry was restored in the OMC. These results suggest that the linkers connect the periplasmic and cytoplasmic domains and stabilize the complex. This is consistent with previous cryo-ET data showing that the linker domain is only present after association of the IMC (*13*).

### Domain Architecture of the System

To obtain a structure of the complete T4SS, we performed a series of focused refinements. First we carried out focused refinement on the previously well-characterized periplasmic region (*8, 10, 12*). The resulting structure at ~21 Å resolution (Fig. S3) matched previously-published *in situ* subtomogram averages well in that region (*8, 10, 12*). The only notable difference compared with other subtomogram averages of that region was the presence of “elbow” density connecting the OMC and periplasmic ring (PR), which is likely due to the presence of sfGFP at the C-terminus of DotF. The collar domain appears to be connected to both the PR and elbow (Fig. 1A and S4). In *Helicobacter pylori*, the collar is also connected to the PR (Fig. S4C). Park et al. (*13*) however suggested that the Dot/Icm T4SS collar connects directly to the OMC through a contact near the DotF-corresponding density (Fig. S4D).

**Figure 1.**
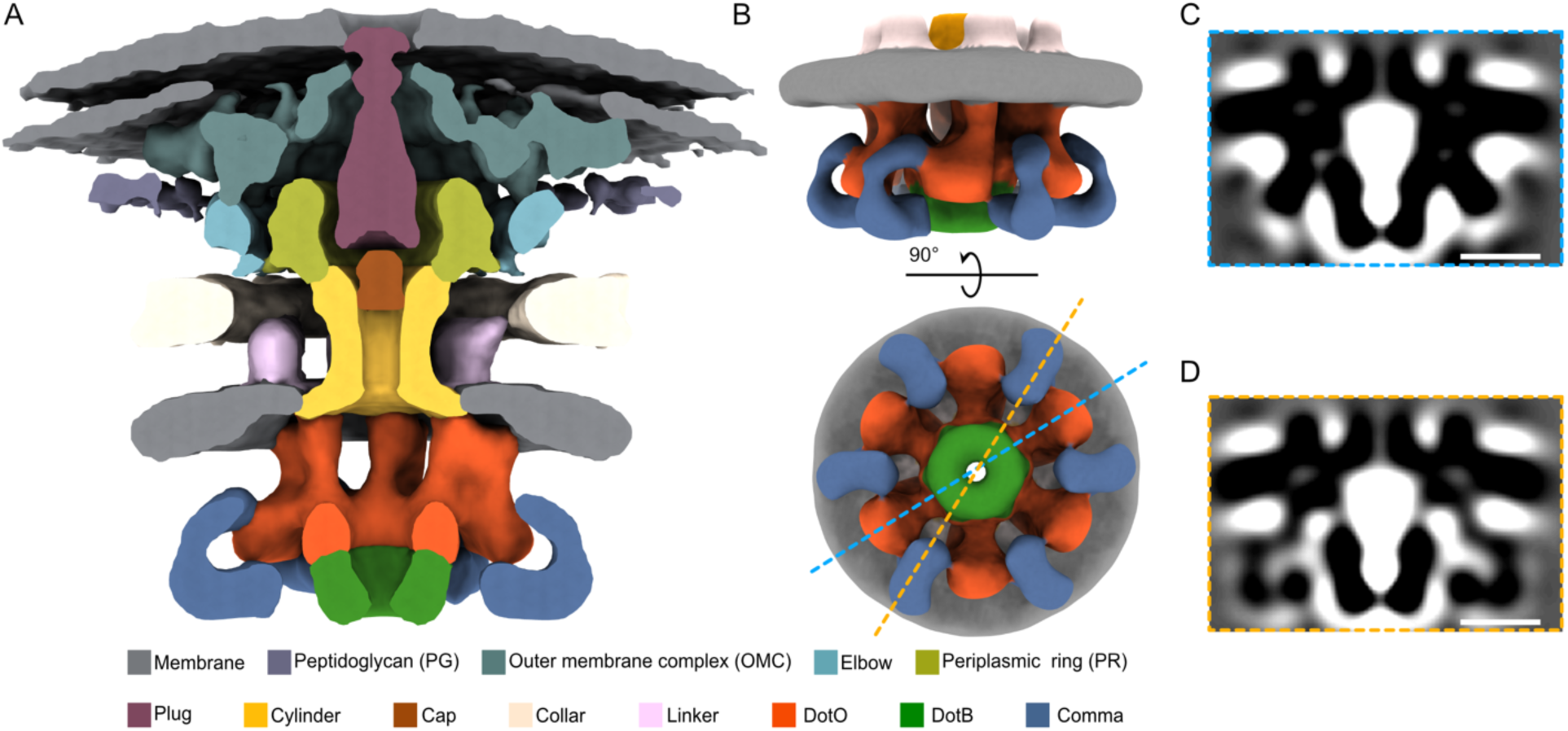
Architecture of the Dot/Icm T4SS. (**A**) Segmented side view of the composite cryo-ET map. (**B**) 3D rendering of the density from focused refinement on the IMC from Class III after applying 6-fold symmetry during post-processing highlighting the novel “comma” density (blue). Densities outside the IMC are masked out. (**C,D**) Central tomographic slices through the symmetrized IMC reconstruction as indicated by corresponding, color-coded, dashed lines in **B**. Scale bars, 10 nm.

Second, we performed focused refinement on the region containing the collar, linker, and a portion of IMC (Fig. S5). A characteristic feature of the T4SS and related systems are symmetry mismatches between individual components. The best characterized mismatches are in the OMC and PR (*15, 16*). More recently, another symmetry mismatch was proposed between the collar and linker domain of Dot/Icm T4SS (*13*). In our average, we observe the transition from a 13-fold symmetry in the collar to 6-fold symmetry in the linker (Fig. S5B-C). The linker domain emerges from the IM and bends towards the collar density (Fig. S5A). Our ability to resolve different symmetries within the same refinement suggests that the elements may be structurally linked.

Third, we performed focused refinement on the IMC in Class III and noticed previously undescribed additional density between the outer ATPases (Fig. 1B-D). In our initial 3D classification, we did not apply any symmetry and the initial alignment used particles with randomized orientations along the symmetry axis of the particle. Even in the asymmetric reconstruction, the uncharacterized density is highly regular (Fig. S6A). To obtain a higher-quality average, we applied 6-fold symmetry to the map (Fig. 1B-D). The uncharacterized density emerges near the bottom of the outside DotO protomers and bends downward in the shape of a comma. While this density has not been described before, we observed a weaker but related structure in the prior map of the IMC from the Liu group (EMD-7612) (*12*) (Fig. S6B-C). In that map, however, the comma density disappears quickly with increasing threshold whereas it does not in our map. One possible explanation is that the presence of the DotF-sfGFP fusion stabilized complete particles and/or allowed better classification (*10*). Finally we merged the maps, using the best-resolved detail at each position, to produce the composite cryo-ET map of the system (Fig. 1A and S1).

### Integrative Model of the Dot/Icm T4SS

The improved quality of our cryo-ET maps, newly-available high-resolution structures of some Dot/Icm T4SS components (*16*), and the recent structure of a related conjugative T4SS (*17*), combined with advances in protein structure prediction (*21*) (Fig. S7) offered us the opportunity to update and extend the molecular model of the *L*. *pneumophila* Dot/Icm T4SS (Fig. 2 and Movie S1). We began by fitting previously known and characterized components of the T4SS. First, we docked the single-particle reconstruction of the OMC and PR (*16*) into the corresponding densities in our cryo-ET map (Fig. S8), accounting for nine components of the system: DotC, DotD, DotF, DotG, DotH, DotK, Dis1, Dis2, and Dis3. As mentioned above, the elbow is most likely a result of the DotF-sfGFP fusion. Consistent with this prediction, we were able to fit 26-copies of sfGFP into the elbow density, resulting in a ring sandwiched by the DotF copies facing in opposite directions (Fig. S9). In the single-particle reconstruction, only the C-terminus of DotF was resolved in both OMC and PR (*16*). Since DotF is known to self-interact (*22*), we built homodimers in ColabFold. We predict that the N-terminal domains of DotF dimerize, forming a coiled-coil structure, and then penetrate the IM with the transmembrane domain, with potentially flexible N-termini exposed in the cytoplasm (Fig. S9D). We show this detail in Fig. S9D, but because corresponding density is not seen in our cryo-ET map, allowing us to place these structures confidently (likely because of flexibility), we did not include the N-terminal domains of DotF in the deposited model shown in Fig. 2.

**Figure 2.**
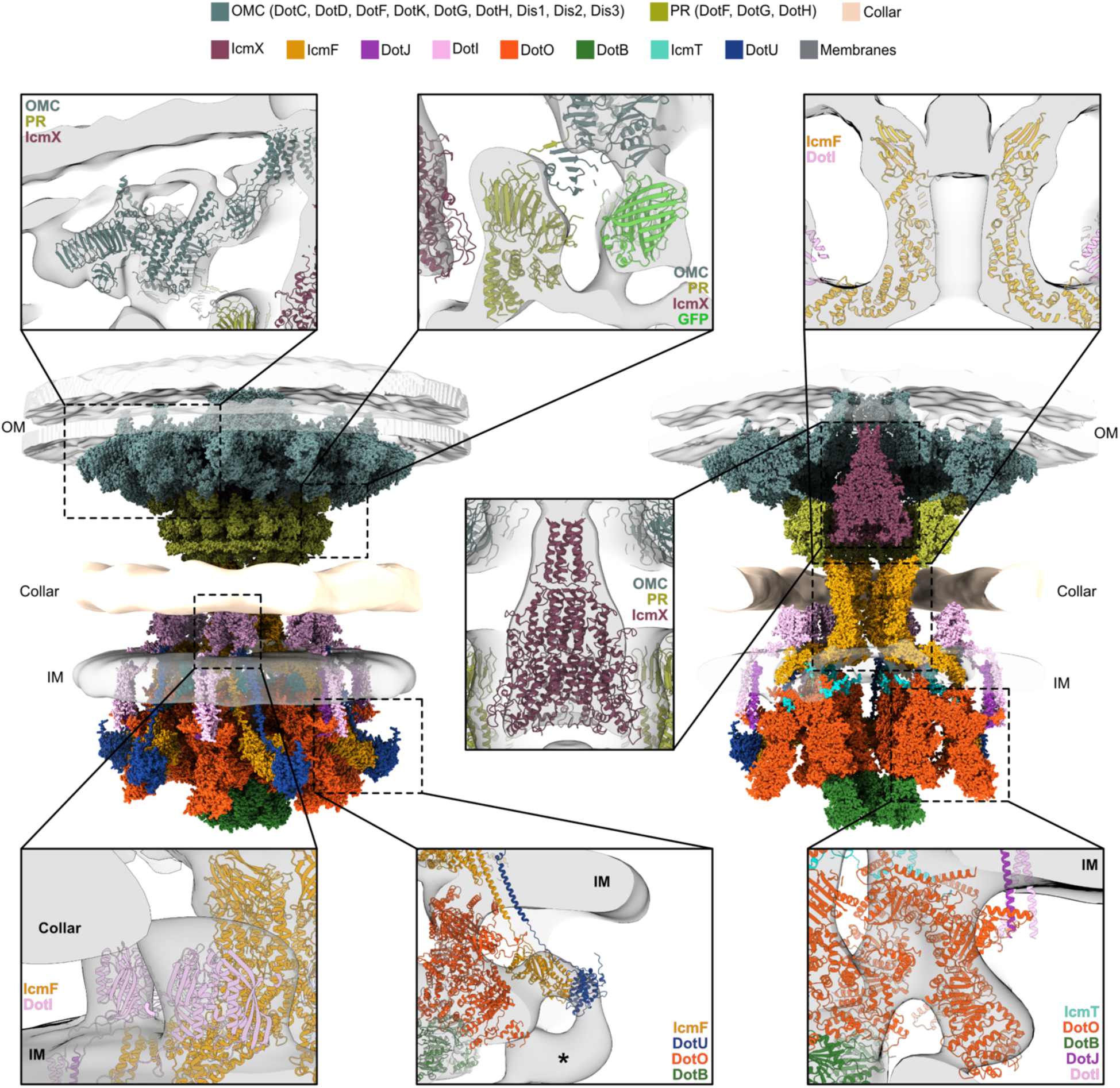
Molecular model of *L*. *pneumophila* Dot/Icm T4SS. Integrative model constructed by docking experimental and predicted protein structures into the *in situ* cryo-ET density maps. The OMC and PR structures are from PDB 7MUS (*16*), while the DotB structure is from PDB 6GEB (*32*). The remaining atomic models are AlphaFold/ColabFold (*21, 23*) predictions. OM - outer membrane; IM - inner membrane; * - unmodeled portion of the uncharacterized comma density in the IMC. The OM, IM, and collar are illustrated as isosurfaces.

Previously, we identified IcmX as a major component of the plug density by comparing difference maps between wild-type and various T4SS mutants (*10*). We ran structure predictions for several different oligomeric states of IcmX, with the best results for a tetramer or pentamer (Fig. S10A-C). The IcmX pentamer had a higher modeling score, as determined by ColabFold (*23*), and occupied a larger volume fraction in our density map. Additionally, we noticed a similarity in shape between the IcmX plug density and the stalk density from the conjugative T4SS (*17*) (Fig. S10D). Notably, the stalk density is generated by VirB5, which is also a pentamer. In addition, IcmX and VirB5 occupy similar “on-axis-in-the-periplasm” positions in the Dot/Icm and conjugative T4SSs, respectively, further indicating their structural and functional relationship. Taking all of this into account, we modeled IcmX as a pentamer into our final model (Fig. 2 and S10E-F).

Between the PR and the IM there is a hollow cylinder. In our previous research, we proposed that DotG may be a crucial component of this cylinder (*10*). No cylinder was seen, however, in the single-particle reconstructions of the purified OMC and PR complex/complexes, which clearly contained DotG (*15, 16*). Later cryo-ET data showing different assembly stages of the Dot/Icm T4SS revealed one stage where the OMC, with a fully-formed dome structure consisting of DotG also had no signs of a cylinder (*13*). Moreover, the N-terminus of DotG is formed by a variable number of 10 amino-acid repeating sequences (Fig. S11A) (*24*). The location of the repeats in DotG, along with their variable number, suggests that DotG does not form the cylinder. Instead, we believe it extends from the PR to form the collar and then penetrates the IM (Fig. S11B). We show this detail in Fig. S11, but the predicted structure for the N-terminus of DotG is low confidence, so we did not dock it into the collar region of our cryo-ET map.

Two additional candidates that could form the cylinder are DotA and IcmF. Given that DotA is a predicted polytopic membrane protein, and assembles into ~10 nm hollow rings upon secretion by planktonic bacteria (*25*), it is unlikely to be a major component of the cylinder. IcmF shares sequence homology with TssM, which forms a decameric (pentamer of dimers) channel within the type VI secretion system (T6SS) (Fig. S12A-B) (*10*). The AlphaFold prediction for the structure of the IcmF monomeric structure has three domains: a cytoplasmic domain, a transmembrane domain, and a periplasmic domain (Fig. S12C-D). The transmembrane domain for IcmF was not annotated in UniProtKB, but DeepTMHMM (*26*) confirmed its presence (Fig. S12C). Six subunits of the C-terminal domain of IcmF fit well into the cylinder density of our cryo-ET map (Fig. 2 and S12E-F).

In T6SSs, TssM (homolog of IcmF) and TssL (homolog of DotU) interact and function to recruit T6SS baseplate proteins to the T6SS membrane complex (*27–29*). In the T4SS DotU and IcmF are co-dependent on each other for stability (*30*) and ColabFold’s IcmF_cyto_:DotU dimer model predicts the proteins also interact within their transmembrane helices and cytosolic portions (Fig. S13A-E). Modeling in these transmembrane domains, the IcmF_cyto_:DotU complex is well-positioned to occupy the upper portion of the comma density (Fig. 2 and S13F).

The IMC is primarily composed of two ATPases, DotO and DotB. DotO is a homolog of VirB4 and when associated with the T4SS complex exists in a unique hexamer of dimer formation at the inner membrane (*12, 17*). DotB is a homolog of VirB11 and is recruited to the DotO dodecamer in an ATP-dependent manner (*19*). Since DotO does not possess a transmembrane domain, it likely requires another protein to recruit it to the membrane. In the conjugative T4SS, VirB4 is anchored to the IM via interactions with VirB3 and VirB8. Six VirB3 molecules bind to the inner six subunits of VirB4, whereas trimers of VirB8 interact with the outer six subunits of VirB4 (*17*). VirB8 trimers further extend to the periplasm via a transmembrane helix where they form periplasmic arches, stabilized by interactions of the C-terminal domain (*17*). In the Dot/Icm T4SS, we found a similarity between IcmT and VirB3 (Fig. S14A). DotI/DotJ were previously recognized as homologs of VirB8 (Fig. S15A) (*10*).

Unfortunately, the resolution of our map is not sufficient to model a protein as small as IcmT on its own. As a result, we built a model of the DotO:IcmT subcomplex with ColabFold and fit it to the density map (Fig. S14). The predicted model shares high structural similarity with the homologous structure from the conjugative T4SS. The completed model of the inner DotO:IcmT hexamer fits the cryo-ET map well (Fig. 2 and S14J). In the conjugative T4SS, each VirB8 trimer creates a periplasmic arch with its N-terminal helices and then elongates a cytoplasmic domain that binds the VirB4 ATPase (Fig. S15B) (*17*). Since biochemical data indicates DotI and DotJ exist as a heterotrimer with a 2:1 DotI:DotJ stoichiometry (*31*), we first built a model of a DotI:DotJ heterotrimer (Fig. S15). We then used it to model the DotI:DotJ:DotO subcomplex, thus allowing us to optimize its position within the cryo-ET map. Finally, we positioned the crystal structure of DotB (*32*) beneath the inner DotO hexamer (Fig. 2 and S16), guided by structural and biochemical data (*13, 19*).

Several transmembrane domains of the Dot/Icm proteins appear to be artificially located below the IM in our final model (Fig. S17A). In our cryo-ET map of class III (fully assembled complex) (Fig. S2A), the IM appears to be less curved than in many individual particles (Fig. S2B). It is likely that the discrepancy we observed is due to variability in the curvature of the IM among individual particles, resulting in the flattening and blurring of the IM in the subtomogram average. As a result, the leaflets of the bilayer are not resolved (as they are in the OM) (Fig. S17A). As we have previously noted, the curvature of the IM is correlated with the resolvability of the linker domain. To investigate this further, we evaluated the effect of the linker-forming proteins (DotI:DotJ) on the membrane curvature prediction. In their absence, the predicted membrane is flat (Fig. S17B), similar to the class I and II averages (Fig. S2A). However, the addition of DotI:DotJ (Fig. S17C) results in a membrane curvature resembling that of the class III average (Fig. S2A). These results are consistent with changes in the IM curvature being associated with the docking of the DotO ATPase via the DotI:DotJ linker.

### Functional adaptations of the IcmF/TssM protein family

TssM has been extensively studied in the context of T6SSs (*33, 34*). Analysis of the cytoplasmic domain revealed the presence of nucleotide binding motifs, and biochemical data confirmed nucleotide hydrolysis activity (*27, 30*). Further detailed sequence analysis revealed that IcmF/TssM belongs to the P-loop NTPase family and contains Walker A and Walker B motifs, typically associated with GTPase activity (*27*). By searching for structural homologs of IcmF_cyto_, we discovered a remarkable similarity with eukaryotic small GTP-binding proteins (Ras) (Fig. 3A-C), a class of proteins that act as molecular switches to quickly turn on and off signaling pathways. Specifically, the AlphaFold model of IcmF_cyto_ shares strong similarity with Arf1, a small G protein that regulates membrane traffic in eukaryotic cells (*35*). Notably, the homology includes highly conserved crucial residues involved in magnesium and GDP binding (Fig. 3B-C). Given this similarity, combined with the observation that TssM is capable of hydrolyzing GTP (*27*), we propose that the cytoplasmic domain of IcmF/TssM proteins contain a Ras-like domain.

**Figure 3.**
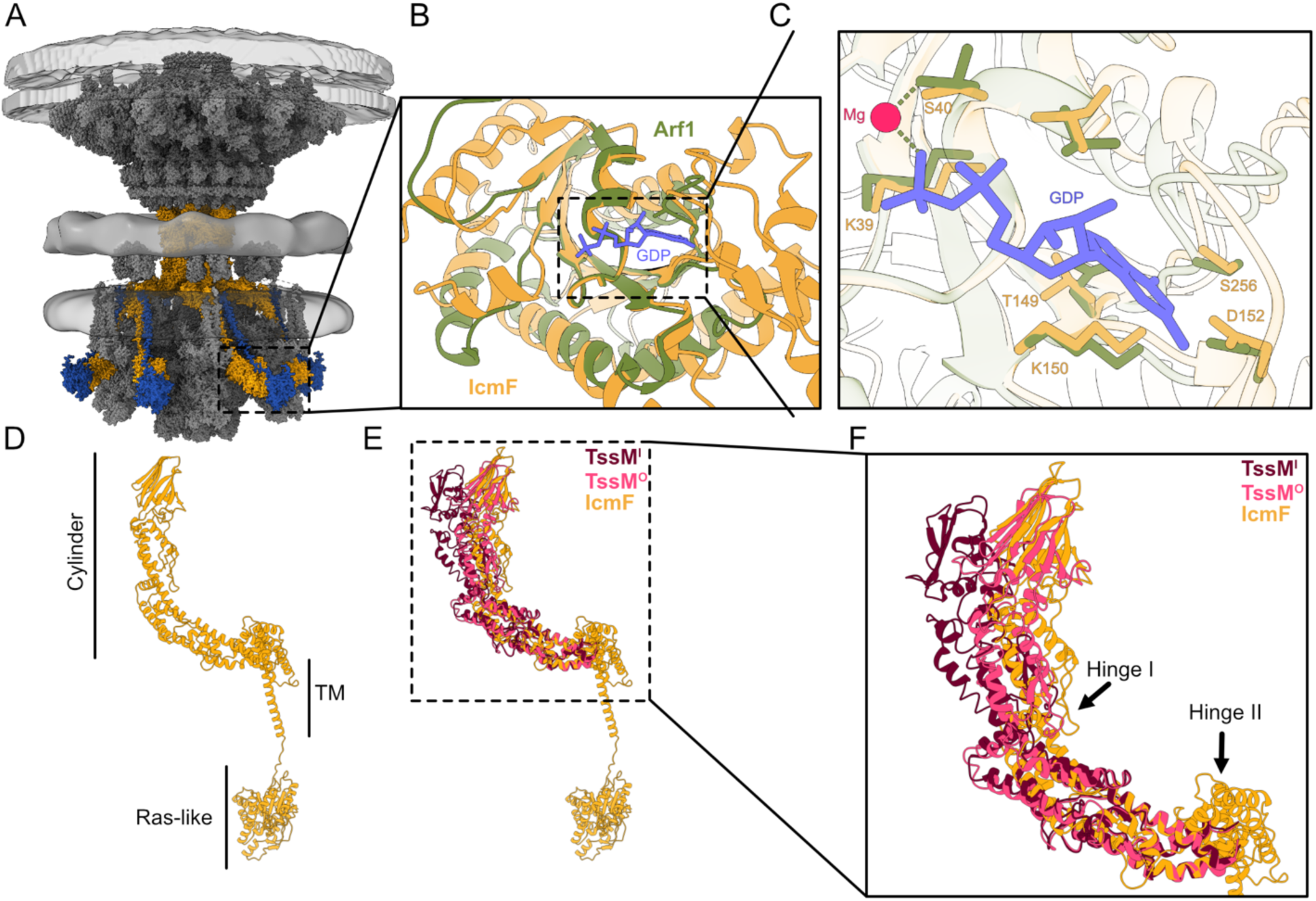
IcmF is a flexible protein with a Ras-like domain. (**A**) Composite model of the T4SS with IcmF and DotU highlighted. (**B**) Cytoplasmic domain of IcmF, which is superimposed with the crystal structure of eukaryotic small G-protein Arf1 (PDB 1R8Q, shown in green) (*35*). (**C**) Structural preservation of the nucleotide-binding site in the IcmF predicted model. A GDP molecule bound to Arf1 and residues critical to nucleotide binding are shown in sticks following the same coloring scheme. (**D**) IcmF domain annotation. (**E**) Structure alignment of IcmF model with cryo-EM structures of TssM from T6SS in two conformations. TssM^I^ and TssM^O^ denote molecules forming the inner and outer parts of the T6SS core complex channel, respectively. (**F**) Enlarged view of the cylinder/channel-forming portion of the IcmF and TssM indicating potential hinge regions.

Another potentially crucial structural adaptation of IcmF is the presence of hinge regions. In the cryo-EM structure of TssM, a hinge region was identified in the middle section of the channel, allowing the protein to rotate and adopt two distinct conformations (*33, 36*). A second hinge region was proposed at the base of the periplasmic domain of TssM (*33*). Both regions are conserved in IcmF (Fig. 3D-F and S12A). IcmF flexibility might play a crucial role in substrate transport through the cylinder and/or contribute to the small cap-like density observed on top of the cylinder in our cryo-ET map (Fig. 1).

### T4SS tethers to the host membrane during early infection

Our atomic model is based on inactive T4SSs expressed in planktonic bacterial cells. We therefore went on to investigate the architecture of actively secreting *L. pneumophila* T4SSs within infected hosts. For host cells, we used human U937 monocytic cells, as they serve as an established model system for the study of *L. pneumophila* macrophage interaction (*34*). In order to slow substrate translocation, and potentially stabilize actively secreting Dot/Icm T4SS complexes that contain a substrate, we challenged U937 cells with *L. pneumophila* cells expressing a chimera of the T4SS substrate RalF with a tightly folded version of ubiquitin (V76G). *Legionella* containing vacuoles (LCV) could be observed associating with mitochondria and ER, typical of infections with wild-type cells (Fig. 4 and S18) (*37*). Ninety-five percent of the T4SS particles found (49/52) exhibited a similar conformation as planktonic T4SS. However, 3 particles were observed in a clearly different state with tethers between the bacterial OM and the membrane of the LCV (Fig. 4 and S18). Similar sites were previously reported in *L. pneumophila* infections of amoeba, suggesting the mechanism of translocation is conserved in the mammalian system (*38*). Notably, the IM in the 3 particles with tethers adopted a sharp curvature similar to that seen in class III planktonic T4SS (Fig. S2A-B), suggesting that the IMC is fully assembled when the T4SS is tethered to the LCV. In contrast to the planktonic class III particles, additional density could be observed between the OMC and PR of the three particles with tethers (Fig. 4C and S18), likely representing T4SS substrates.

**Figure 4.**
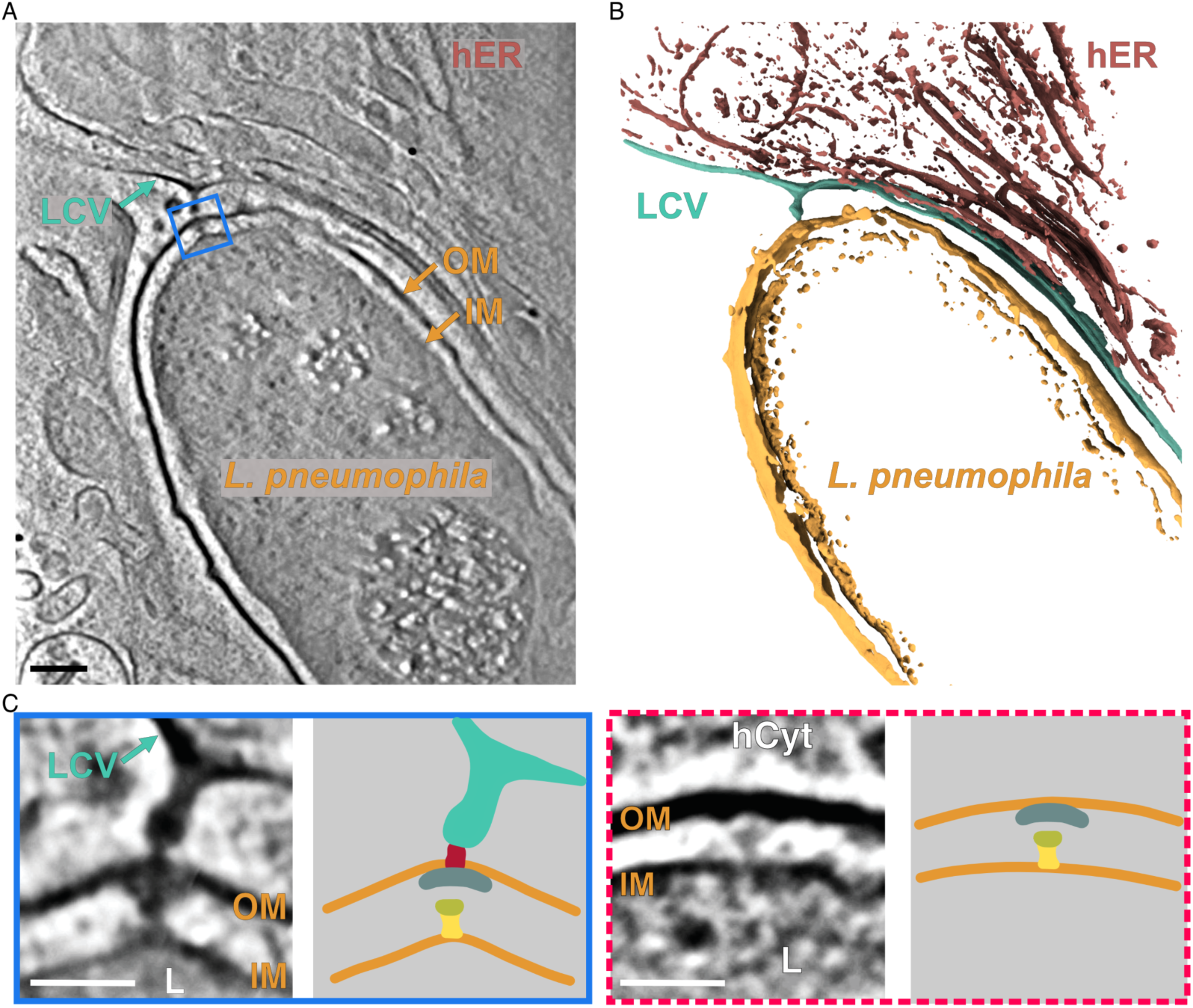
*L. pneumophila* T4SS tethers to the host membrane during early infection. Representative tomographic slice (**A**) and segmented model (**B**) illustrating a T4SS observed inside *L. pneumophila* cells infecting human U937 cells at 1 h post-infection. OM (orange) - *L. pneumophila* outer membrane; IM (orange) - *L. pneumophila* inner membrane; LCV (cyan) - *L. pneumophila* containing vacuoles. L (orange) - *L. pneumophila* cells; hER (brown) - Endoplasmic reticulum of U937 cells. Scale bar, 100nm. (**C**) Enlarged T4SS particles identified in A (blue frame) and a different cell shown in Fig. S17C (red dashed frame). The particle tethered to the host membrane is enclosed by the blue frame; the particle without tethering site is indicated by the red dashed frame. The panels on the right depict the individual particles making the OMC, PR, and the cylinder schematically. Scale bars, 50nm.

## Discussion

The *L. pneumophila* Dot/Icm T4SS is a large macromolecular machine composed of ~30 proteins. Twenty-two of the proteins form the core system and 8 are part of a distinct membrane-associated complex, the Type IV Coupling Protein (T4CP) complex (*39–41*). Of the 22 proteins forming the core system, our model contains 17 proteins, with over 460,000 total atoms, comprising one of the largest and most complex macromolecular structures modeled to date. Compared to the structure of the conjugative T4SS published last year (*17*), our model revealed four major novelties, which are discussed individually below (Fig. 5).

**Figure 5.**
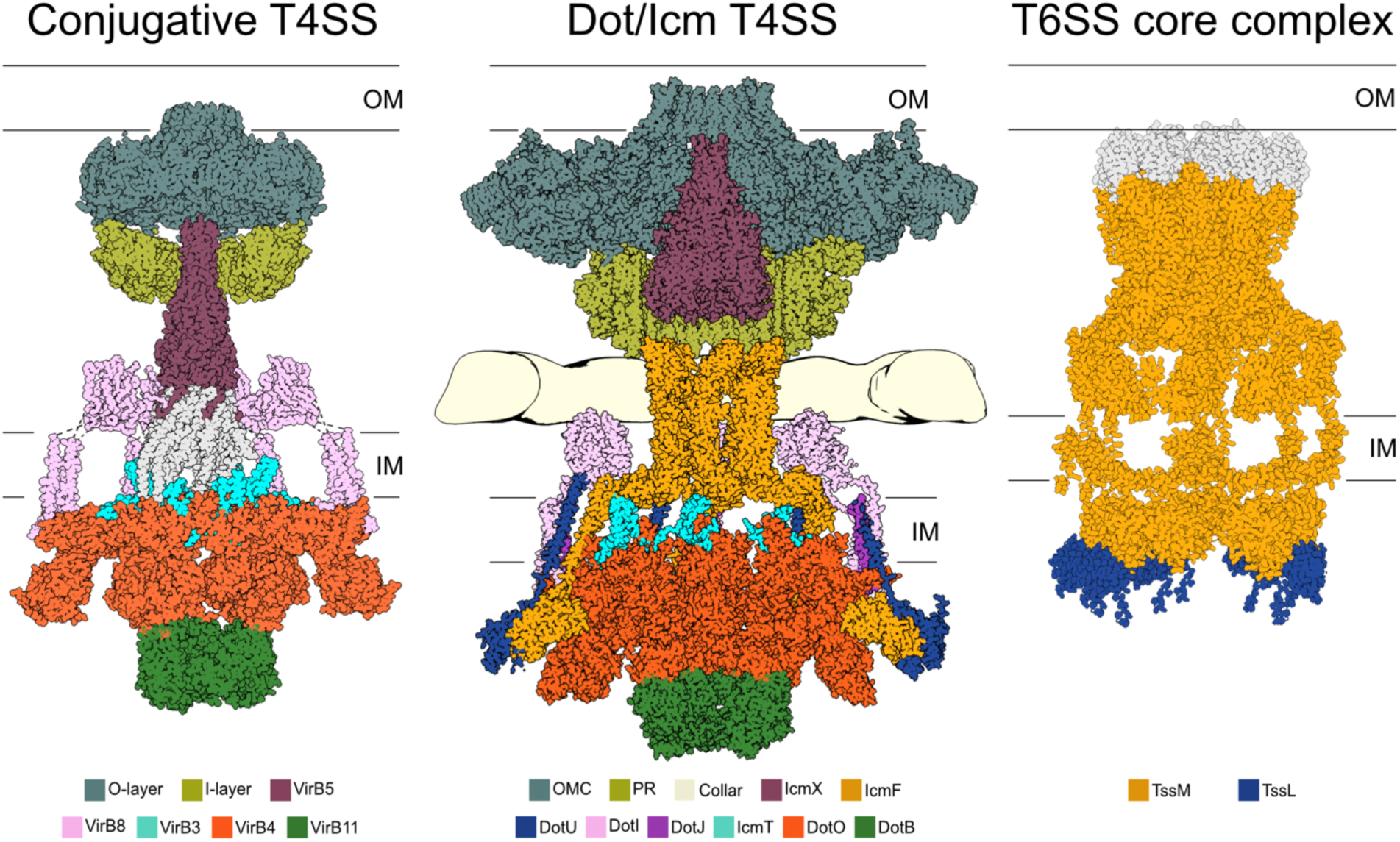
*L. pneumophila* Dot/Icm T4SS is a hybrid system. Left panel - atomic model of the conjugative T4SS (PBD 7O3J, 7O3T, 7O3V, 7O41) (*17*) with VirB11 model (PDB 2GZA, shown in green) (*86*) placed as proposed by Mace et al. (*17*). Central panel - integrative model of the *L. pneumophila* Dot/Icm T4SS (this study). Right panel - composite model of T6SS core complex. Periplasmic domain (TssJ/TssM) is from the single particle cryo-EM reconstruction (PDB 6IXH) (*34*), the cytosolic domain of TssM (yellow) is an AlphaFold prediction, and TssL is from a crystal structure (PDB 3U66) (*36*). Cytosolic domains of T6SS are placed based on our previous structure of the T6SS (EMD-8600) (*87*). Homologous components between the systems are marked with the same colors.

### IcmF cylinder

The first difference is the presence of IcmF forming a central hollow cylinder in the Dot/Icm T4SS in place of the conjugative T4SS component VirB6. In the conjugative system, VirB6 forms a pentameric, conical platform on the axis of the system adjacent to the IM (*17*). Mace et al. (*17*) proposed that VirB6 scaffolds the assembly of the pilus, which is made up of VirB2 subunits. In contrast, the Dot/Icm system does not appear to encode a VirB6 homolog or any pilin subunits, nor has a pilus structure been observed. Although the Dot/Icm T4SS appears to have been derived from an IncI plasmid conjugation system, additional adaptations were apparently required to secrete the wider variety of substrates (over 300) exported by *L. pneumophila* into host cells (*24, 42*). These changes included the incorporation of new components such as secretion chaperones and DotU/IcmF, which are homologs of the T6SS factors TssL/TssM (*36, 43*). Since the Dot/Icm system does not contain a VirB6 homolog, and the periplasmic portion of IcmF is located in a similar place as VirB6 in the conjugative system (Fig. 5), we propose the VirB6 platform in conjugative T4SSs was replaced with IcmF in the Dot/Icm T4SS. In T6SSs, TssM forms a channel just above the IM through which the Hcp rod of the T6SS (a large cargo) is projected (*44*); in the Dot/Icm T4SS, IcmF appears to form a central hollow cylinder through which substrates traverse from the IMC to more distal portions of the T4SS complex in the periplasm. In addition, both IcmF and TssM contain hinge regions, which may be critical to the export of larger sized cargo.

### Comma density, IcmF_cyto_ and GTP hydrolysis

The second major difference observed in our model is the presence of a cytoplasmic density which we called the comma. We assigned the upper portion of this density to the cytoplasmic domains of IcmF and DotU. TssL/TssM, homologs of DotU/IcmF, function to stabilize the T6SS complex and recruit an assembly platform of proteins known as the baseplate (*27–29*). Similarly, DotU/IcmF stabilize the T4SS complex as part of their role in mediating optimal Dot/Icm substrate export (*30*). In contrast to TssL/TssM, however, DotU/Icm perform an additional function as they are necessary and sufficient to target the Dot/Icm T4SS to the bacterial poles (*10*). Based on the interaction of TssL/TssM’s cytoplasmic domain with other T6SS baseplate components, it is likely that DotU/IcmF’s cytoplasmic domains bind another factor(s) and this may account for the density in the lower portion of the comma. Potential candidates for this factor(s) include a polar-localizing determinant, an as-yet-unidentified Dot/Icm component, and/or possibly an early-translocated Dot/Icm substrate.

We further found that IcmF_cyto_ consists of a Ras-like domain, but the role of GTP hydrolysis remains unclear. One possibility is that nucleotide hydrolysis by IcmF_cyto_ is required for the proper targeting and/or assembly of the Dot/Icm T4SS complex. Alternatively, nucleotide hydrolysis could mediate conformational changes in the cytoplasmic ATPases and cylinder, allowing substrates to engage with the complex and/or cause the IcmF cylinder to open. Finally, secretion through the T4SS is contact-dependent (*45–47*), and our *in situ* data shows T4SS particles making contact with the host membrane. Another possibility is therefore that host contact induces a signal that is transmitted through the periplasmic components of the Dot/Icm T4SS to IcmF_cyto_, resulting in GTP hydrolysis and recruitment of other factors to the complex.

### Collar

The third major difference is the presence of the periplasmic density we called the collar. We propose that a portion of the N-terminus and/or the middle domain of DotG containing the repeats forms the collar for several reasons. First, in the *Helicobacter pylori* Cag T4SS, the resolved fragment of CagY, the DotG homolog, in the PR faces the collar, which hints that the unresolved repeating fragment could extend further to form the collar (*48*). This is further suggested by a well-resolved connection between the PR and collar in our earlier cryo-ET structure of the Cag T4SS (*9*). Second, we note that there is no similar repeat-rich, middle domain of VirB10, the DotG homolog in conjugative systems, and no collar has been seen in structures of those systems (*17, 49, 50*). Finally, there are 13 copies of DotG in the OMC, and we found that the collar also has 13-fold symmetry. Unfortunately, the low resolution of the collar density and the lack of confident structural predictions for these domains of DotG prevented us from modeling this region. The purpose of the collar remains unclear. One intriguing possibility is that the repeat-rich middle domain of DotG allows the T4SS cylinder to expand in either width or height to accommodate large conformational changes during substrate export or penetrating a host cell membrane.

### Plug

The fourth major difference is the plug, composed of IcmX in the Dot/Icm system. Interestingly, both IcmX and VirB5 form pentameric cone-shaped structures on-axis in the periplasm. VirB5 is later found at the tip of extended pili (*17*), however, and the Dot/Icm system is not thought to build a pilus. The IcmX pentamer is also located closer to the OM than VirB5. Other differences include the Dot/Icm T4SS being only partially assembled in an Δ*icmX* mutant, with no cylinder and an altered conformation of the PR (*10*), and IcmX being released from planktonically grown bacteria (*51, 52*). Based on these observations, we hypothesize that IcmX functions either as a scaffold for assembly of the Dot/Icm T4SS and/or as a plug in non-secreting T4SSs, waiting to be expelled at the onset of substrate export.

### DotI:DotJ linkers

One similarity between the conjugative and Dot/Icm systems is the presence of a hexameric set of linkers spanning the IM. Previously, we reported that the linker/collar region (formerly referred to as “wings”) appears flexible (*10*). The linker formed by DotI:DotJ complexes provides a docking site at the IM for the outer six subunits of the DotO dodecamer. Based on their location, another potential role of DotI:DotJ could be sensing conformational changes from the outer DotO subunits during substrate transport, propagating them through the collar to the OMC/PR, thereby allowing for plug release and secretion of effector proteins.

### Secretion pathway

While here we have emphasized the novel role of IcmF in forming a hollow cylinder through which substrates pass, the entire Dot/Icm system appears to be an extended channel. Effectors likely first interact with the T4CP DotL ATPase, then pass through the opening of the dodecameric DotO ATPase, across the IM, through IcmF and finally the PR/OMC. Although data suggest that unfolded T4SS substrates are preferred (*41, 53*), we could not identify any obvious hydrophobic patches along the secretion path that would protect hydrophobic fragments of unfolded targets from refolding. Considering the wide variety of effector proteins secreted through the T4SS (*40*), it is possible that some substrates are secreted in folded or partially-refolded forms. In support of that notion, expulsion of IcmX would create a chamber of sufficient size to allow small proteins to refold.

### Integrative modeling

Much of cell biology is driven by large, multi-subunit flexible complexes that are challenging or impossible to purify or reconstitute, so may only be available for study within intact cells. Here we have demonstrated an integrative approach to characterize such structures. New cryo-ET instrumentation, methods, and software (*20, 54–56*) made it possible to image the T4SS directly *in situ* (*57*) and overcome stoichiometric variance by collecting and classifying thousands of particles. Focused refinement mitigated the effects of flexibility. Recent developments in single-particle cryo-EM methods (*15, 16*) and structure prediction with AlphaFold (*21, 23, 58*) provided atomic models of individual components and sub-complexes. The docking of these structures into our cryo-ET maps illustrates what is becoming possible in a new era of integrative modeling.

## MATERIALS AND METHODS

### *L. pneumophila* strains preparation

*L. pneumophila* strain (JV9082) expressing a DotF-sfGFP from the chromosome was constructed by allelic replacement into strain Lp02 using the suicide plasmid pJB7255 (DotF-sfGFP). pJB7255 was constructed by cloning two PCR products (generated by PCR using primers JVP2973/JVP2990 and JVP2992/JVP2993, respectively) into the R6K suicide plasmid pSR47S (Table S1). The PCR products were digested with BamHI/NotI and NotI/SacI, respectively, and ligated into BamHI/SacI-digested pSR47S.

*L. pneumophila* strain (JV9298) expressing DotB(E191K) and DotF-sfGFP from the chromosome was constructed by allelic replacement into strain JV9082 using the suicide plasmid pJB7487. This plasmid was constructed by subcloning a ~1.2 kb fragment (BamHI/SalI) containing DotB (E191K) from plasmid pJB2920 (*19*) into BamHI/SalI-digested pSR47S.

### Sample preparation for cryo-ET

*L. pneumophila* strain (JV9082 and JV9298) was cultured as previously described (*8*). In short, *L. pneumophila* were grown to stationary phase in CYE media supplemented with 150 ug/mL ferric nitrate, 400ug/mL of L-cysteine and 100ug/mL of thymidine. Bacteria cells were collected and resuspended in the same growth media to OD_600_ = 6.0. Resuspended culture was mixed with 10 nm gold beads as fiducial markers and then applied onto glow-discharged copper R2/2 200 mesh Quantifoil holey carbon grids (Electron Microscopy Sciences) in Vitrobot Mark IV (Thermo Fisher Scientific). Grids were plunge frozen with liquid ethane/propane mixture (*59*) and stored at liquid nitrogen temperature.

### Cryo-ET data collection

Tilt series were collected at 300 keV at pixel sizes 2.65 Å/pix on a Titan Krios (Thermo Fisher) equipped with a K2 or K3 camera. One-hundred micron objective aperture was inserted during collection and a 20 eV slit energy filter (GATAN EFTEM) was used to improve contrast. For whole cell tomograms, either cell pole was targeted for collection. Tilt series were collected bi-directionally from 0 deg to 60 deg on each side, with 2 or 3 deg steps. A total of 100 electrons/Å^2^ was distributed evenly across the tilt series. Tilt images were collected either as a single image or a stack of 10 frames in the super resolution mode.

### Tomograms reconstruction and subtomogram averaging

Raw image stacks were motion corrected with MotionCor2 (*60*) and assembled into tilt series with the IMOD newstack program (*61*). For subtomogram averaging in EMAN2, tilt series alignment and subsequent tomogram reconstruction was performed automatically within the EMAN2 tomography pipeline (*62*). EMAN2 reconstructed 4* binned tomograms were used in the IMOD 3dmod program to manually pick particles. An effort was made to include as many top views as possible. Particle coordinates were picked to be centered visually at the point where the cylinder meets the IM; an orientation was also recorded to have the particles aligned as closely as possible to what is shown in Fig. 1A. The center and orientational information were then converted to particle records in EMAN2. Subsequent subtomogram averaging was performed using the manual alignment information as the starting point in EMAN2. We used 2* binned particles for the initial round of averaging, focusing on the density in the periplasmic space with C13 symmetry enforced. Then aligned particles were allowed to move locally with a cylindrical mask containing the whole particle. The alignment information was used to extract unbinned particles and the same local refinement was repeated. Next, sub-tilt refinement was performed to obtain the best StA map for the whole particle.

For subtomogram averaging in Dynamo (*63*) and I3 (*20*), tilt series alignment was performed either manually or automatically in Etomo on 2* binned tilt images. We used tomo3d (*64*) to reconstruct 2* binned weighted back-projection tomograms from the aligned tilt series. Then we used an in-house script to transfer the aligned particle coordinates into a Dynamo compatible form (script available: https://doi.org/10.5281/zenodo.7659617) and performed StA in Dynamo with a mask enclosing the entire particle and C13 symmetry enforced. The aligned particles were extracted again from the tomograms. These sub-tomograms were treated as individual tomograms containing one particle in I3. The particle alignment orientation was also transferred into I3 with an in-house script (script available: https://doi.org/10.5281/zenodo.7679253). Initially I3 StA was carried using a soft-edge cylindrical mask enclosing the entire system to exclude particles without assembled IMC using classification. The rotational angle around the C13 symmetry axis of the remaining particles were randomized and the particles were further classified using a soft-edge cylindrical mask around the IMC. The class with the fully assembled IMC complex was used for further focus refinement on either the IMC or connection between the cytoplasm and periplasm. At any step of data processing in I3 we did not apply symmetry to the reference map. Final reconstructions with well-resolved symmetry were eventually symmetrized to aid model building. Crucial steps in the data processing pipeline are summarized in Fig. S1. The resolution estimation is performed on the unfiltered, unsymmetrized maps using ResMap (*65*) (Fig. S3). The composite map was created in ChimeraX using *vop maximum* command (*66*).

### Subtomogram averaging - data analysis

To test symmetry of the collar, the cryo-ET map from focused refinement on collar/IM connection was further masked out to only include the collar ring. Rotational cross-correlation was calculated using *dynamo_symmetry_scan* function in Dynamo (*63*) (Fig. S5B).

Using an in-house python script (script available: https://doi.org/10.5281/zenodo.7679388), particle shift between two iterations was calculated as the total shift of the particle center as recorded in the I3 output after each iteration.

### Cryo-FIB milling

*L. pneumophila* strain (JV9298) was cultured similarly as for the whole cell tomography. Grid preparation follows the same procedure as described above, except that bacterial cells were resuspended to OD_600_ = 20.0 and applied onto glow-discharged copper R2/2 200 mesh Quantifoil holey carbon grids. Excess material on the grid was removed by manual blotting from the back before plunge-freezing. Frozen grids were mounted into Autogrids (Thermo Fisher Scientific) and transferred into an FEI Versa 3D equipped with a Quorum PP3010T Cryo-FIB/SEM preparation system (Quorum Technologies LLC, East Sussex, UK) operating at cryogenic temperatures. Samples were sputter coated with a thin layer of platinum (15 mA, 60 sec) and then imaged with the SEM at 10 keV. Identified areas with multiple layers of bacterial cells were further subject to stepwise milling with 30keV gallium ions, starting from 0.3 nA beam current and gradually stepping down to 10 pA, to generate final lamellae approximately 150-300 nm thick (*67*). Tomograms were reconstructed in IMOD (*68*) and processed with IsoNet (*69*).

### Infection assay for cryo-ET

Human U937 cells (ATCC CRL-1593.2) were grown in RPMI1640 supplemented with 10% FBS and L-glutamine for several passages and then subjected to a 72-h incubation with 12-O-tetradecanoylphorbol-13-acetate (TPA) for differentiation. Differentiated U937 cells became adhesive and were seeded onto gold London Finder grids (Au H2 R2/2, Quantifoil) coated with vitronectin (25 µg/mL in 1xPBS, pH 7.4), allowing for a 48 h incubation at 37°C. *L. pneumophila* Lp02 expressing CyaA tagged RalF attached to an irremovable ubiquitin chain (Lp02 CyaA-RalF::V76G) (*53*) was induced at OD_600_ = 2 to 2.5 with 250 µM IPTG for 4-5 h. U937 cells were infected at an MOI of 100 for either Lp02 CyaA-RalF::V76G or JV9082 and vitrified after 1 h in the presence of 10 nm gold beads as fiducials. Tilt-series were collected bi-directionally at 33 000 x equipped with the Volta Phase Plate from −54° to 54° with an increment of 3°. A total of 100 electrons/A^2^ was distributed evenly across the tilt-series. Each tilt-image was collected in the super resolution mode either as a single image or a stack of frames. Tilt-series were phase corrected and reconstructed with SIRT in IMOD (*60*). Tomograms were further processed with IsoNet (*69*) and manually segmented with Amira (ThermoFisher).

### Modelling

All the maps were segmented using the Segger package in ChimeraX (*66, 70, 71*). Available experimental structures for OMC/PR (*16*) and DotB (*32*) were fitted in the map using the *fitmap* command. The atomic models of the remaining components were predicted using AlphaFold/ColabFold (Fig. S7) (*21, 23, 58*). We used the R388 T4SS structure (PDB 7OIU) (*17*) as scaffold for the positioning of DotO, IcmT and DotJ:DotI_2_ atomic models. Detailed information on placing Dot/Icm T4SS components into our density maps are described below.

### OMC/PR

The single particle cryo-EM of the OMC/PR (*16*) was fitted as a rigid body into the corresponding density in the cryo-ET map from focused refinement on the periplasmic complex (Fig. S8). Fitting was guided by the 13-fold symmetry of the OMC.

### DotF/GFP

DotF consists of a TM in position 53-75 followed by a coiled-coil region (135-176). The C-terminal portion of DotF has been solved (208-269) and is occupying two different positions: a C13 ring beneath the Dis3 protein of the OMC and a C18 ring in the PR (*16*). DotF IM localization and its ability to self-interact has been reported before (*22*). Considering the coiled-coil region, we build a DotF dimer in ColabFold. The model dimerizes in this region with long flexible linkers connecting to the N- and C-terminal. We positioned the coiled-coil region of this model halfway between two DotF C-terminal domain solved structure and the TM helix in density corresponding to the IM. We build the linkers in MODELLER (*72*) and symmetrized it with the respective symmetry of the PR and OMC (Fig. S9).

Given the DotF-sfGFP high local concentration the sfGFP may dimerize or form high molecular weight aggregates (*73*). To our knowledge the only available structure of a native GFP dimer is the extra-superfolder GFP (PDB 5B61) (*74*) a close homolog of the sfGFP superimposing with a RMSD of 1.8 Å (*74*). We fitted the structure in the collar by rigid docking and applied C13 symmetry (Fig. S9).

### IcmX

IcmX as a main component of the plug was identified based on the analysis of the T4SS mustants (*10*). However, symmetry of the IcmX is unknown. For that reason, we tested structure predictions for multiple symmetries as generated by ColabFold. The best results were obtained for tetramer and pentamer with overall scores for pentamer being higher. Thus, we fitted IcmX pentamer as a rigid body into the plug density in ChimeraX (Fig. S10).

### DotG

DotG repetition were identified using the MOTIF search tool of the Japanese GenomeNet service (*75*). As previously reported (*24*) five copies of the Pentapeptide_4 repeat motif was found. Given the low score ranging from 3.1e-06 to 6.8e-12 and the Alphafold DotG model not showing the classic right-handed β-helical structure typical of the pentapeptide repeat motif we investigated the repetition sequence. The GenomeNet identified repetition were aligned in MEGA using MUSCLE (*76, 77*). For this multiple alignment we were able to identify a 10 amino acid pattern, as previously described (*22*), starting with a Lys or Arg and ending with a Leu or Iso. We therefore extracted those sequence and built a sequence logo using WebLogo3 (*78*) to identify the motif sequence. This motif ([KR]-X-A-G-[FY]-[STND]-X-X-X-[LI]) is different form the pentapeptide repeat sequence ([STAV][DN][LF][STR][G]) and explain the discrepancy of the DotG AlphaFold model with the solved structures. In order to investigate the fold of this repetition we scanned the PDB database with this motif but only the DotG protein was found.

### IcmF:DotU

IcmF and DotU proteins are present in both T4SS and T6SS (TssM and TssL, respectively) (*36*). The T6SS proteins have been extensively characterized and the structure of both proteins has been solved (*33, 34, 36, 79–81*). The TssM monomeric structure is characterized by the presence of three domains spanning across the inner membrane. The periplasmic portion consists of four helix-bundle domains connecting a β-sheet rich domain, involved in TssJ binding. The periplasmic domain is connected by a TM helix to the cytoplasmic ATP binding domain (*27*). TssL:TssM interaction involves a 110-aa TssM extension of the nucleotide binding domain (*28*) and a cleft on the TssL structure (Asp74, Glu75, Gly137 and Phe138) (*29*).

We closely inspected IcmF and DotU AlphaFold models for T4SS and T6SS shared features. In the IcmF Uniprot page there is no TM helix annotated, but DeepTMHMM (*26*) predicted, with high confidence, a TM in position 317-338 (Fig. S12C). While a C-terminal TM helix is predicted for DotU. Sequence alignment of DotU with TssL revealed conservation of the residue involved in TssM binding (Fig. S13A). Given the high conservation of IcmF and TssL predicted interacting regions we run AlphaFold multimer. In the IcmF_Cyto_:DotU dimer model the proteins are interacting with the cytosolic portion of IcmF and DotU with the two transmembrane helices in a good orientation. We built the IcmF cylinder by manually placing it in the map region forming the central cylinder and applying C6 symmetry (Fig. 12). We then manually placed the IcmF_cyto_:DotU in the newly identified comma density and manually rebuilt the IcmF linker and reoriented the DotU TM helix to cross the membrane density in our map (Fig. 13).

### DotO:IcmT

The DotO biological assembly consists of DotO dimers arranged in a hexamer (*13, 17*). Interaction between DotO and IcmT allows DotO-membrane tethering. We therefore built a DotO_2_:IcmT trimer atomic model. ColabFold modeled the DotO dimer in the isoform involved in hexamer assembly rather than the dimer formation. Therefore we used R388 T4SS structure (PDB 7OIU) (*17*) as a template to position the outer DotO subunits. Superimposition of the DotO:IcmT with VirB4(TrwK):VirB3(TrwM) revealed a good RMSD between the two structures and similar positioning of IcmT in comparison to TrwM (Fig. S14F). A DotO monomer was further aligned with its VirB4 (TrwK) counterpart to generate the dimer. The DotO_2_:DotT trimer was fit into the map from focused refinement on the IMC using Powerfit (*82*) and C6 symmetrized in ChimeraX. This generated a clash between DotO C-terminal domains in the inner protomers. In the Colabfold DotO dimer this region is predicted to fit in a cleft region of the other subunit (Fig S14G). We used this prediction to remodel the clash.

### DotI:DotJ

To model the DotI:DotJ subcomplex we submitted to ColabFold a 2:1 DotI:DotJ assembly based on the available biochemical data (*31*). This model can be divided into two regions: a transmembrane helix bundle and the periplasmic part connected by a flexible loop. The TM helix trimer superimposes well with the VirB8 (TrwG) triple helix (RMSD = 0.501), preserving the position of the TM segment. Therefore, we used this conformation for the positioning of the helix bundle in DotO proximity. The periplasmic portion consists of the two DotI globular domains modeled with the alpha helices facing the interior of the complex. This orientation is the same as one of the conformations resolved in the DotI crystal (Fig. S15G) (*31*). This configuration is also adopted by VirB8 (TrwG) for the formation of the arches region in the conjugative T4SS (*17*). We manually positioned the DotI periplasmic part in the linker density and applied C6 symmetry. The flexible linkers were built using MODELLER (*72*).

### DotB

The DotB structure has been solved and characterized by mutagenesis study. *In vivo* cryo-ET studies revealed its ability to induce a conformational change in DotO upon binding (*13, 19, 32*). We used the available mutation data (*19*) to place DotB into the cryo-ET map below the inner DotO hexamer (Fig. S16).

### Bioinformatics and visualization

Membrane simulation. Since the IM in our STA is bend, we used PPM3.0 web server (*83*) to investigate proteins orientation. This server allows protein-induced membrane deformations using membrane systems reflecting the properties of gram-negative inner membrane.

Protein sequence alignment was performed using Clustal Omega (*84*) and visualized with Jalview (*85*). Data were visualized using GraphPad Prism, IMOD (*61*), and ChimeraX (*66*).

## ACKNOWLEDGEMENTS

Electron microscopy was performed in the Beckman Institute Resource Center for Transmission Electron Microscopy at Caltech and Pacific Northwest Center for Cryo-EM (PNCC). We thank the Liu lab for sharing data and expertise in using I3 to resolve the DotO hexamer of dimers. Y.L. was supported by a Jane Coffin Childs Memorial Fund for Medical Research post-doctoral fellowship. This work was supported by the National Institutes of Health (grant R01-AI127401 to G.J.J.).

## AUTHOR CONTRIBUTIONS

P.D. prepared samples, reconstructed tomograms and picked particles for a subset of data, performed subtomogram averaging, performed data exploration, located proteins forming the system, participated in building the model, drafted the manuscript, and prepared the figures. Y.L. conceived experiments, prepared samples, acquired and reconstructed tomograms, picked particles and worked on subtomogram averaging, performed the particle shift analysis, planned and led FIB-milling and infection experiments, and contributed to writing of the manuscript. S.M. performed data exploration, identified proteins forming the system, led model building, prepared the movie explaining the model building and helped write the manuscript. D.G. conceived experiments, prepared samples, acquired and reconstructed tomograms, and helped interpret the data. J.W. reconstructed tomograms, picked particles for a subset of data, worked on FIB-milling and infection experiments, performed segmentation, participated in data interpretation, and helped to write the manuscript. S.D.C. helped plan and execute the infection experiments. W.Z. helped plan and execute the FIB-milling experiments. S.V. constructed *L. pneumophila* strain JV9298. J.P.V. constructed *L. pneumophila* strain JV9082 and helped revise the final version of the manuscript. G.J.J. supervised the entire project from experimental design through interpreting the results and helped write the manuscript.

## DECLARATION OF INTERESTS

The authors declare no competing interests.

## DATA AVAILABILITY

Cryo-ET maps and Dot/Icm T4SS integrative models will be uploaded to public repositories and made available upon publication. During revision process data are available upon request. Scripts used for data analysis are available on Zenodo: https://doi.org/10.5281/zenodo.7659617, https://doi.org/10.5281/zenodo.7679253, https://doi.org/10.5281/zenodo.7679388).

---

**Figure S1.**
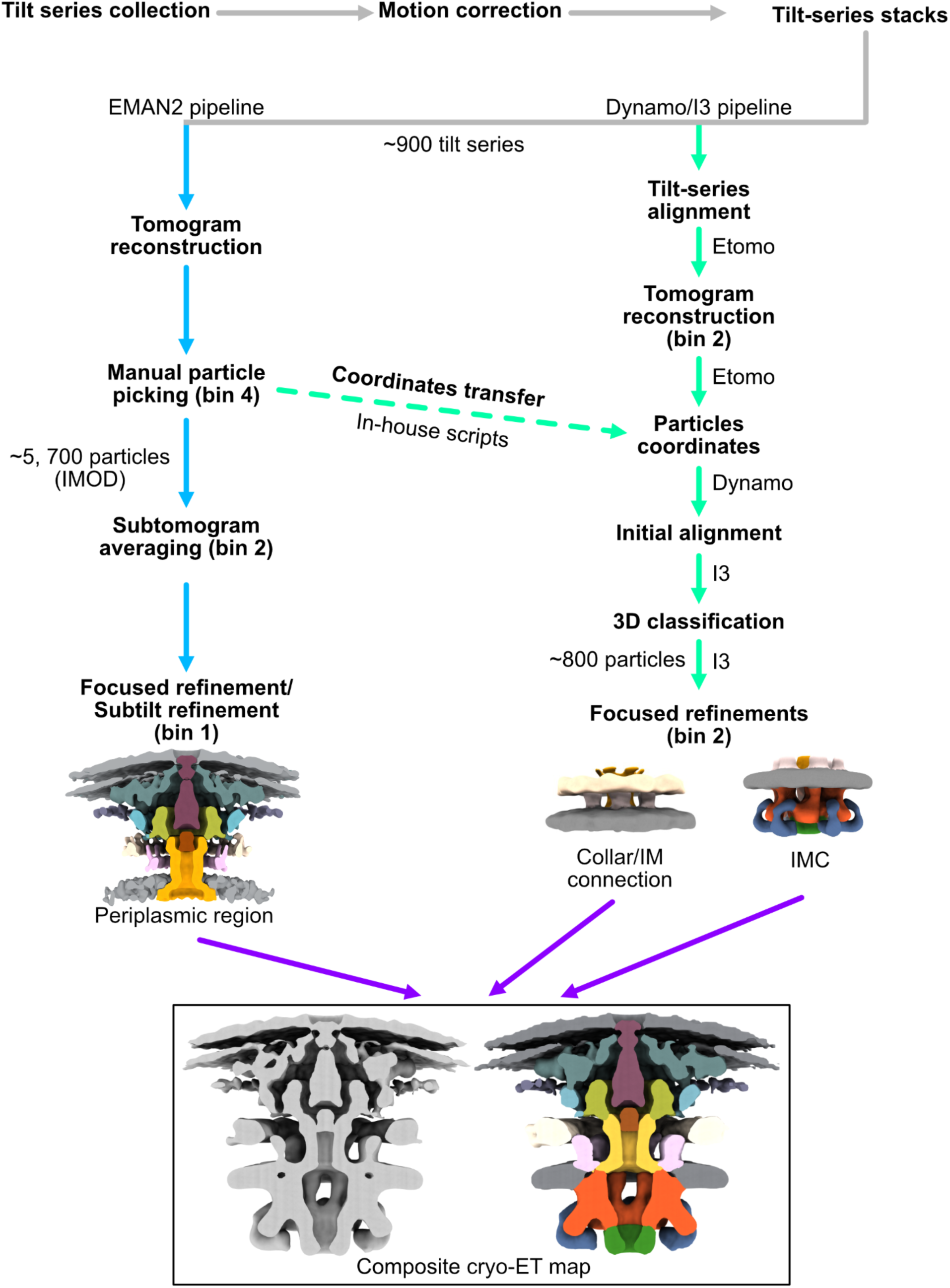
CryoET data processing. Simplified schematic of the data processing workflow highlighting crucial steps in the pipeline.

**Figure S2.**
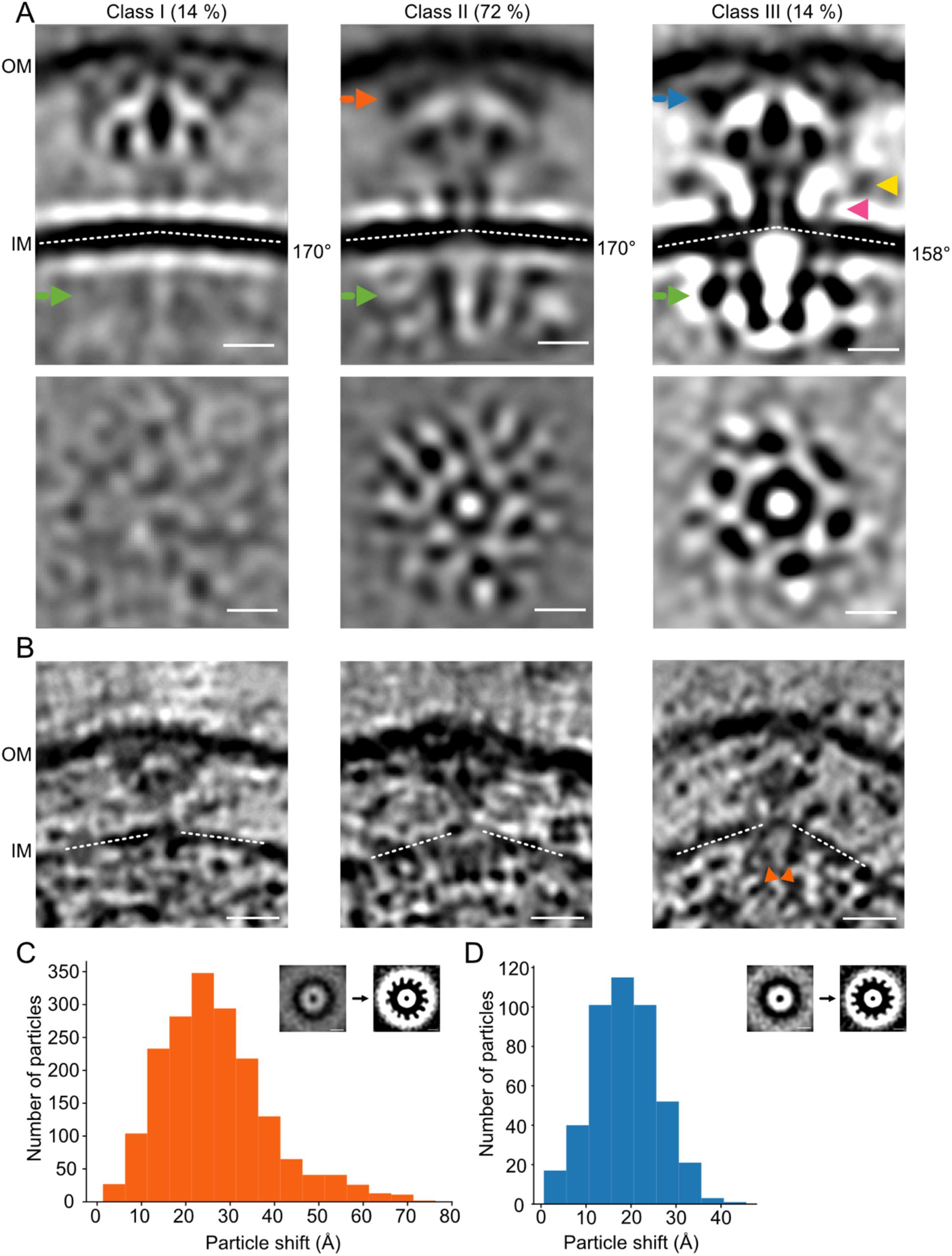
Compositional Heterogeneity of the inner membrane complex (IMC) *in situ.* **(A)** 3D class averages of T4SS with different compositions of the IMC. Top panel, side view. Bottom panel, cross-section at the position indicated by the green arrows in the top panel. Percentages indicate the number of particles within each class. OM - outer membrane; IM - inner membrane. Yellow and pink arrows indicate the collar and linker densities, respectively. Scale bars, 10nm. (**B**) Individual T4SS particles from FIB-milled samples witch matching composition. Scale bars, 20 nm. Orange arrows indicate the cytosolic ATPases. (**A**,**B**) IM curvature is marked by white dashed lines. (**C**,**D**) Histograms showing total shifts required to restore 13-fold symmetry in the OMC for particles included in Class II (**C**) and Class III (**D**) after focused refinement on the IMC. Insets show a cross-section at the position indicated by orange (**C**) and blue (**D**) arrows in (**A)**, respectively. The left and right subpanels represent OMC structure before and after symmetry recovery. Scale bars, 10 nm.

**Figure S3.**
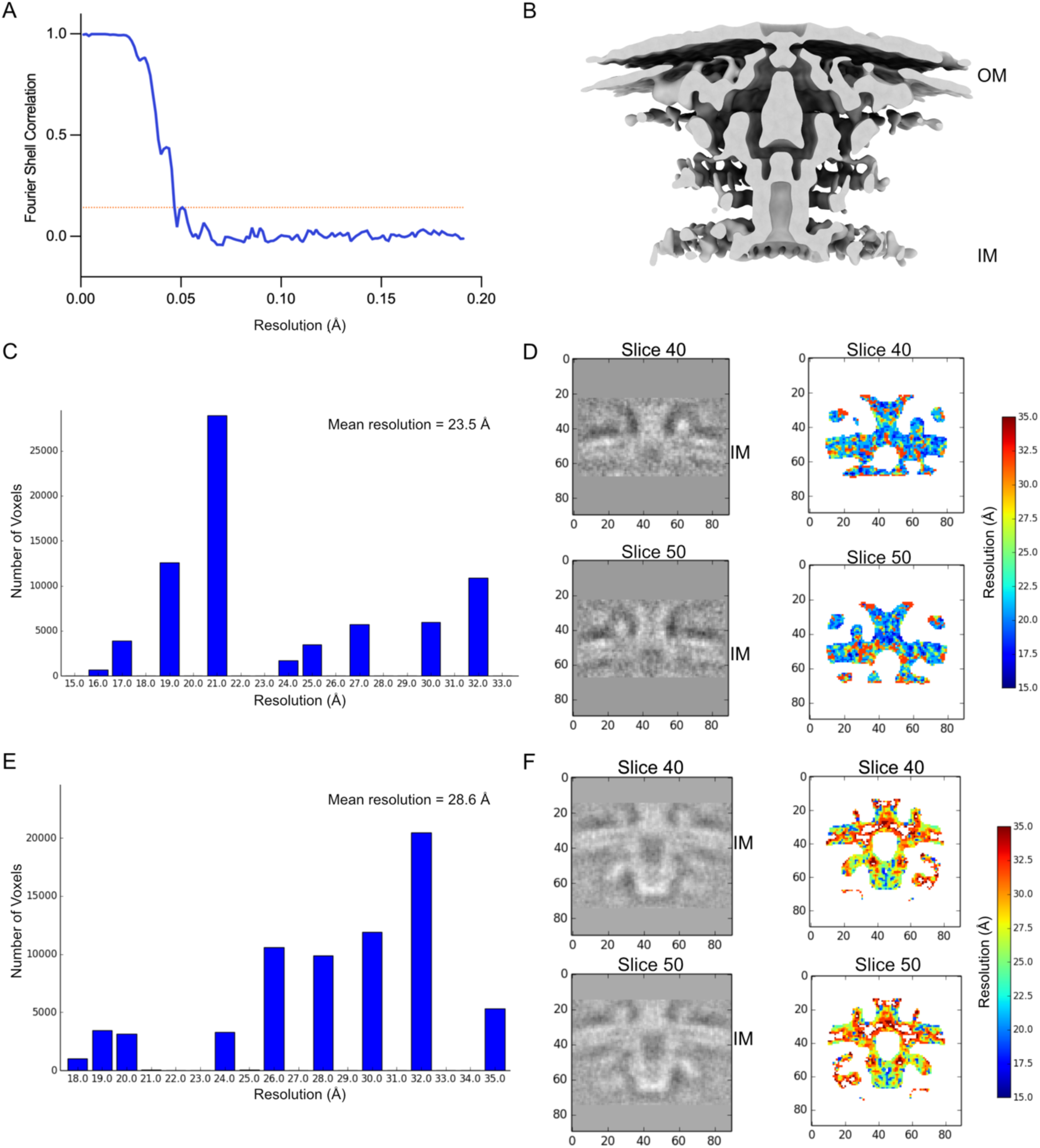
Resolution estimations. (**A**-**B**) Resolution estimation of the EMAN2 subtomogram average of the periplasmic region with C13 symmetry applied. (**A**) Fourier Shell Correlation (FSC) curve with an orange horizontal line indicating a 0.143 threshold. FSC_0.143_ = 21 Å. (**B**) Side view of the corresponding density map. (**C**-**F**) Because gold-standard processing is not implemented in I3, the resolution estimation for maps from I3 was analyzed using ResMap. Resolution was estimated using unfiltered, symmetrized maps. (**C**-**D**) Resolution estimation for the map with local refinement on the collar/IM connection. (**E**-**F**) Resolution estimation for the IMC.

**Figure S4.**
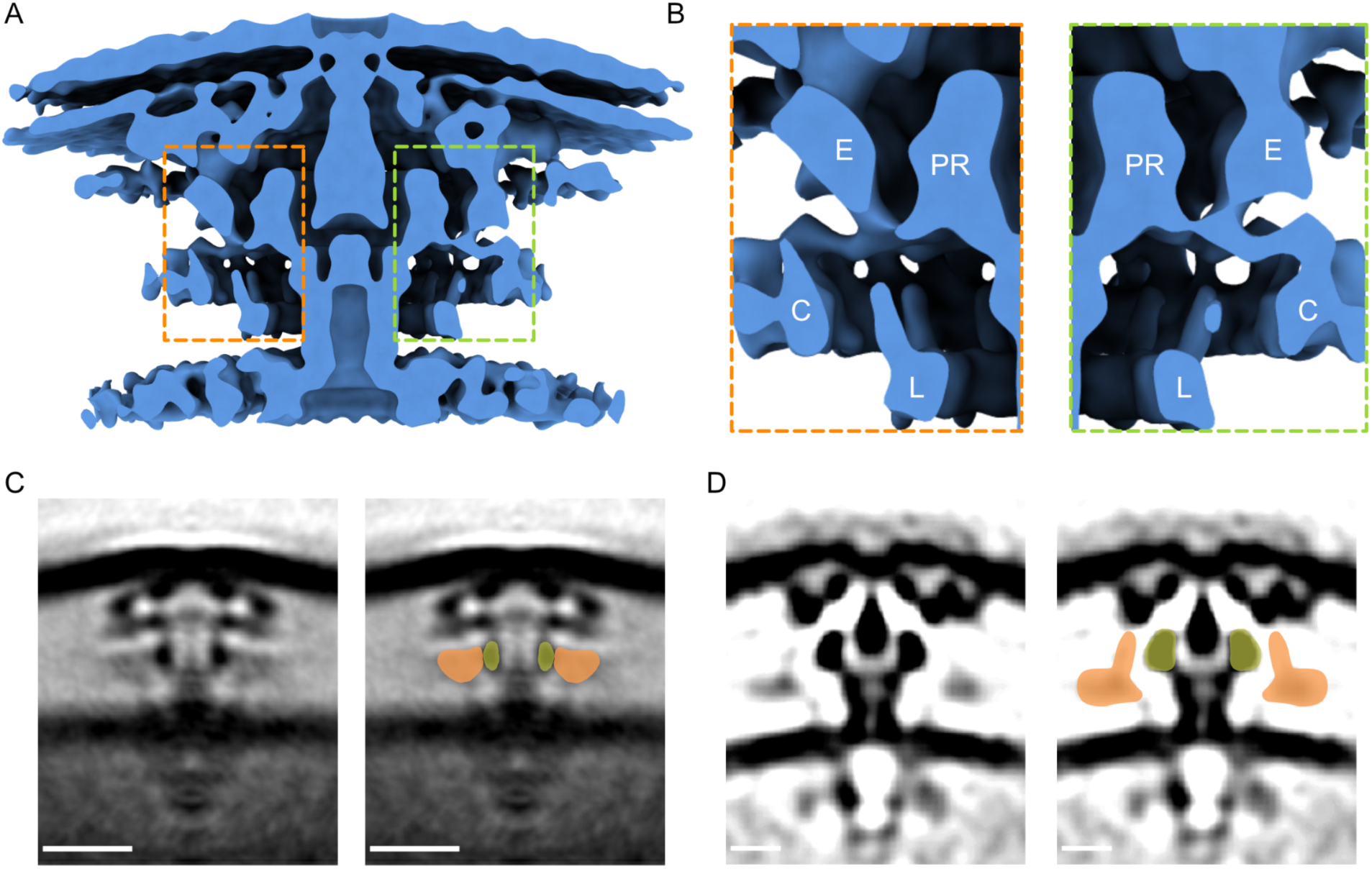
Connectivity of the collar density. (**A**) Side view of the periplasmic complex. (**B**) Enlarged sections as indicated in (**A**) by dashed outlines. E - elbow; PR - periplasmic ring; C - collar; L - linker. (**C**) Central slices through sub-tomogram averages of *H. pylori* cag T4SS (EMD-7474) (*9*). Scale bars, 20 nm. (**D**) Central slices through sub-tomogram averages of *L. pneumophila* Dot/Icm T4SS reproduced from Fig. 2E in Park et al. (*13*). Scale bars, 10 nm. (**C**,**D**) In panels on the right, beige and green marked densities indicate collar and PR, respectively.

**Figure S5.**
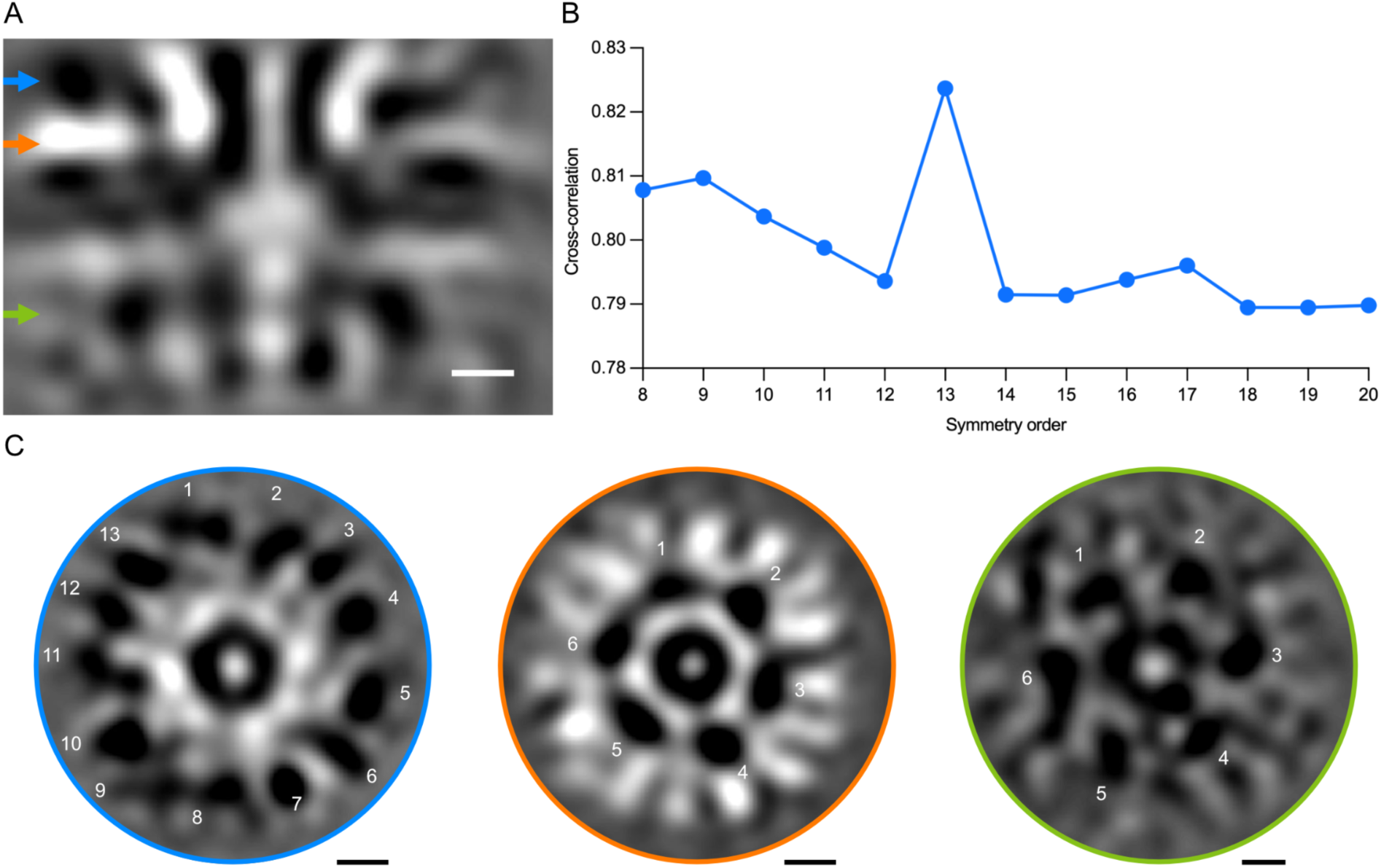
Symmetries of the collar and IMC. (**A**) Central tomographic slice from the focused refinement of the collar/IM region. (**B**) 3D rotational cross-correlation for the collar region marked by a blue arrow in (**A**). (**C**) Cross-sectional views at the positions indicated by color-coded arrows in (**A**). Scale bars, 10 nm.

**Figure S6.**
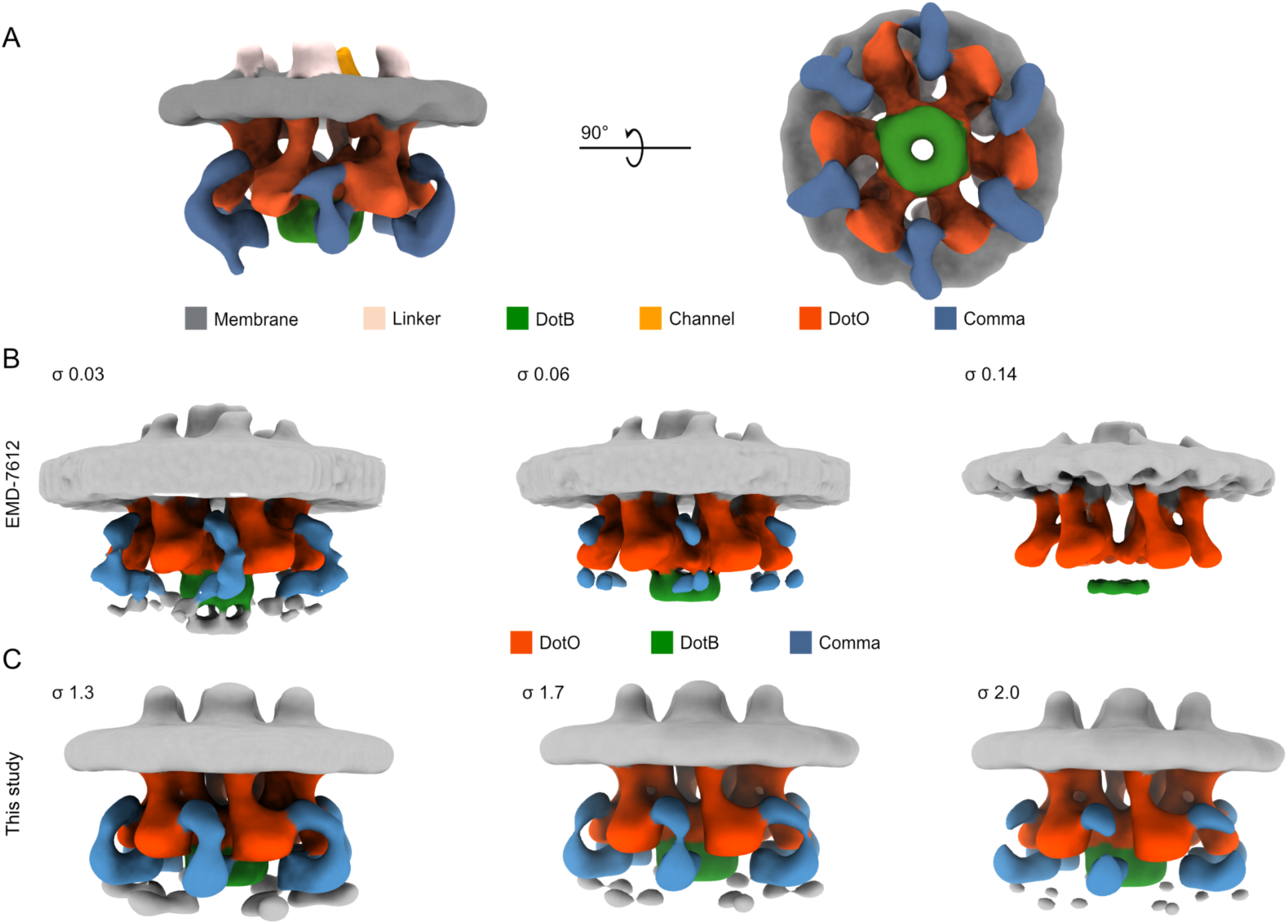
Previously unreported density in the IMC. (**A**) 3D rendering of the unsymmetrized density from focused refinement on the IMC from Class III (Fig. S2A). Densities outside the IMC are masked out. (**B**,**C**) Cryo-ET density maps of the IMC displayed at different threshold levels with colored densities corresponding to DotO, DotB, and uncharacterized comma density. (**B**) EMD-7612 (*12*). (**C**) This study.

**Figure S7.**
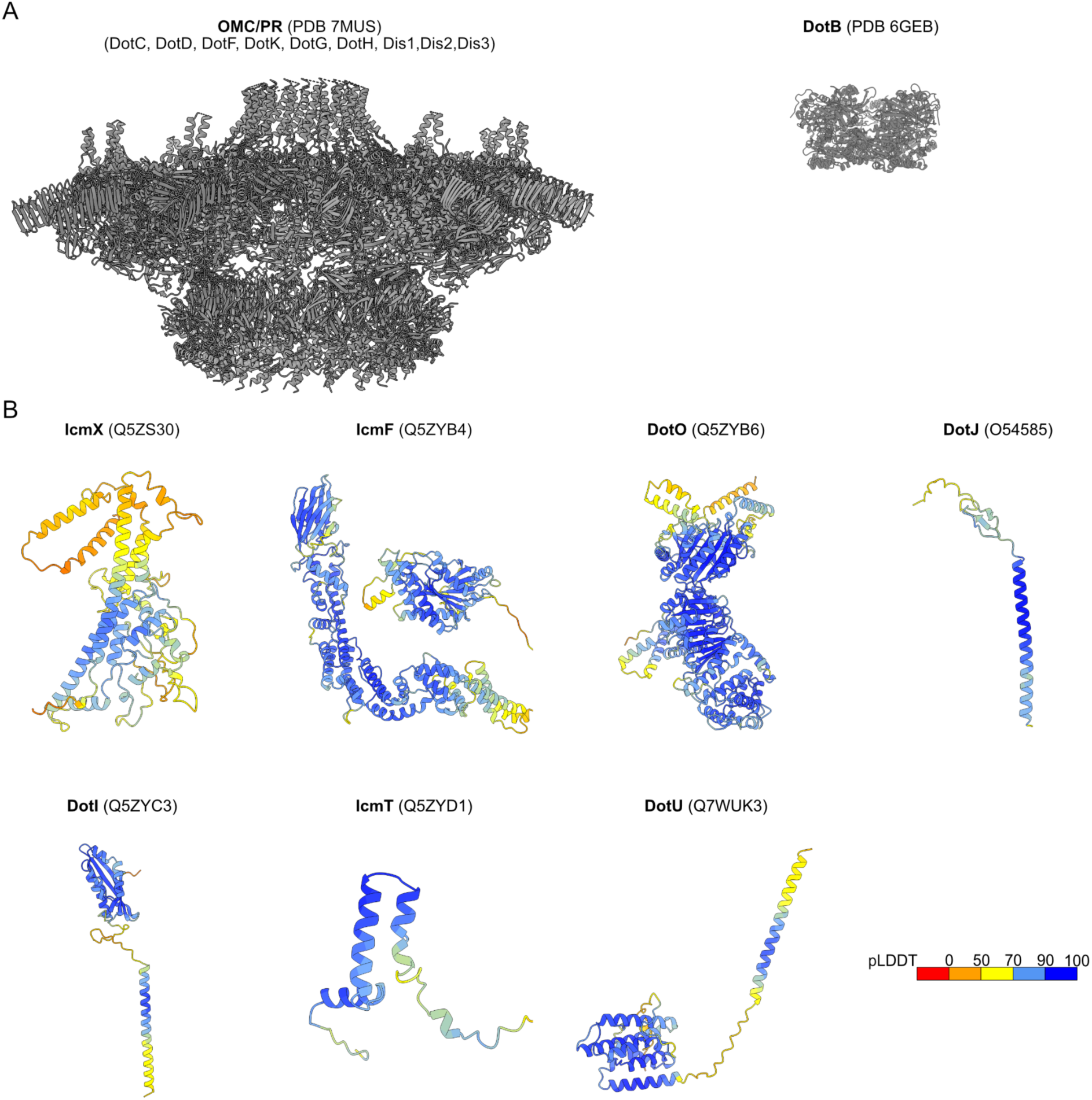
Dot/Icm T4SS components used to build the integrative model. (**A**) Experimentally determined structures. (Left) Single-particle cryo-EM structure (PDB 7MUS) of the periplasmic core complex (*16*) and (right) crystal structure (PDB 6GEB) of DotB (*32*). (**B**) Monomeric AlphaFold structure predictions for Dot/Icm T4SS components. The models are colored by local model confidence estimated with a predicted local distance difference test (pLDDT) (*21*). UniProtKB accession numbers are indicated in the parenthesis.

**Figure S8.**
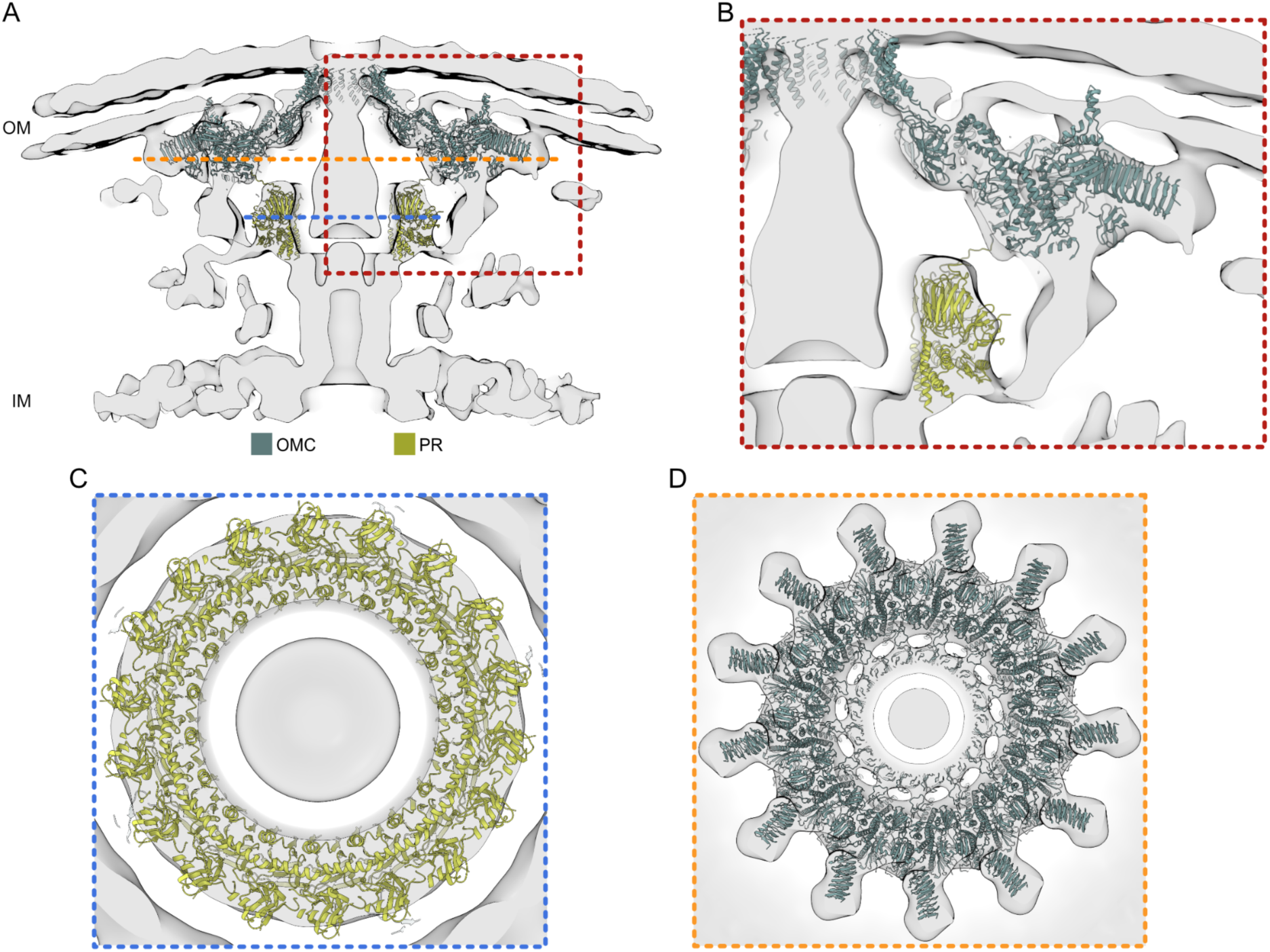
Fitting the atomic models of the OMC and PR into the cryo-ET density map. (**A**) 13-fold symmetrized cryo-ET map (gray) of the T4SS periplasmic region with the atomic models of OMC and PR (PDB 7MUS) (*16*). (**B**) Enlarged section from (**A**) as indicated by the red dashed box. (**C**,**D**) Cross-sections at the positions indicated by blue (**C**) and orange (**D**) dashed lines in panel (**A**).

**Figure S9.**
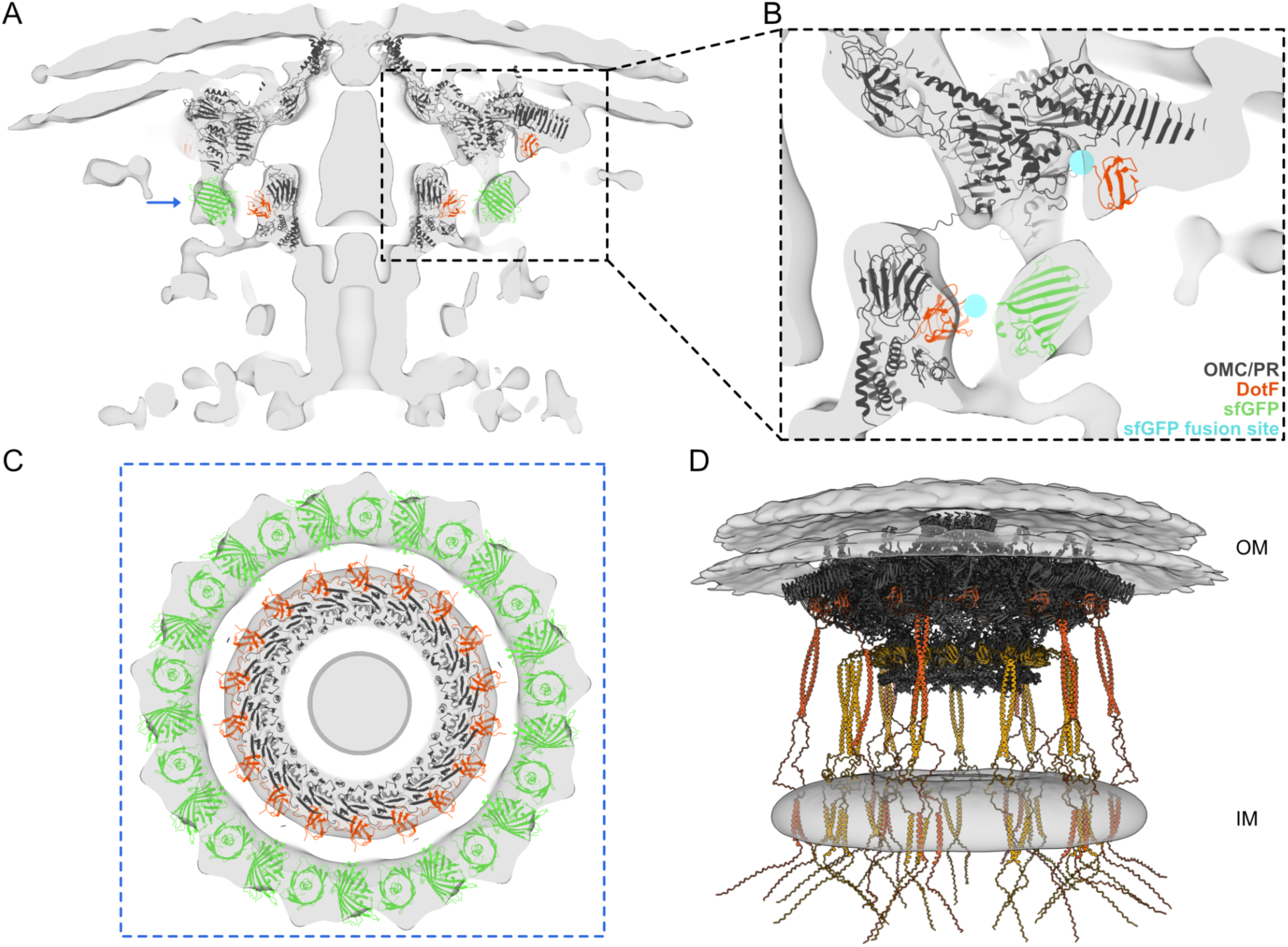
Elbow density attributed to sfGFP. (**A**) Atomic models of the OMC and PR (PDB 7MUS) (*16*) fit into the cryo-ET map. DotF subunits are colored in orange. The atomic model of sfGFP (green) dimer (PDB 5B61) (*74*) fit into the elbow density. (**B**) Inset shows enlarged section of the elbow density. The position of DotF-GFP fusion in our construct is indicated by blue spheres. (**C**) Cross-section the elbow density ring at the positions indicated by a blue arrow in panel **A**, with 26-copies of sfGFP. (D) Predicted localization of the coiled coil and transmembrane N-terminal domains of DotF. Subunits of the DotF portions localized in the OMC and PR are colored in orange and yellow, respectively.

**Figure S10.**
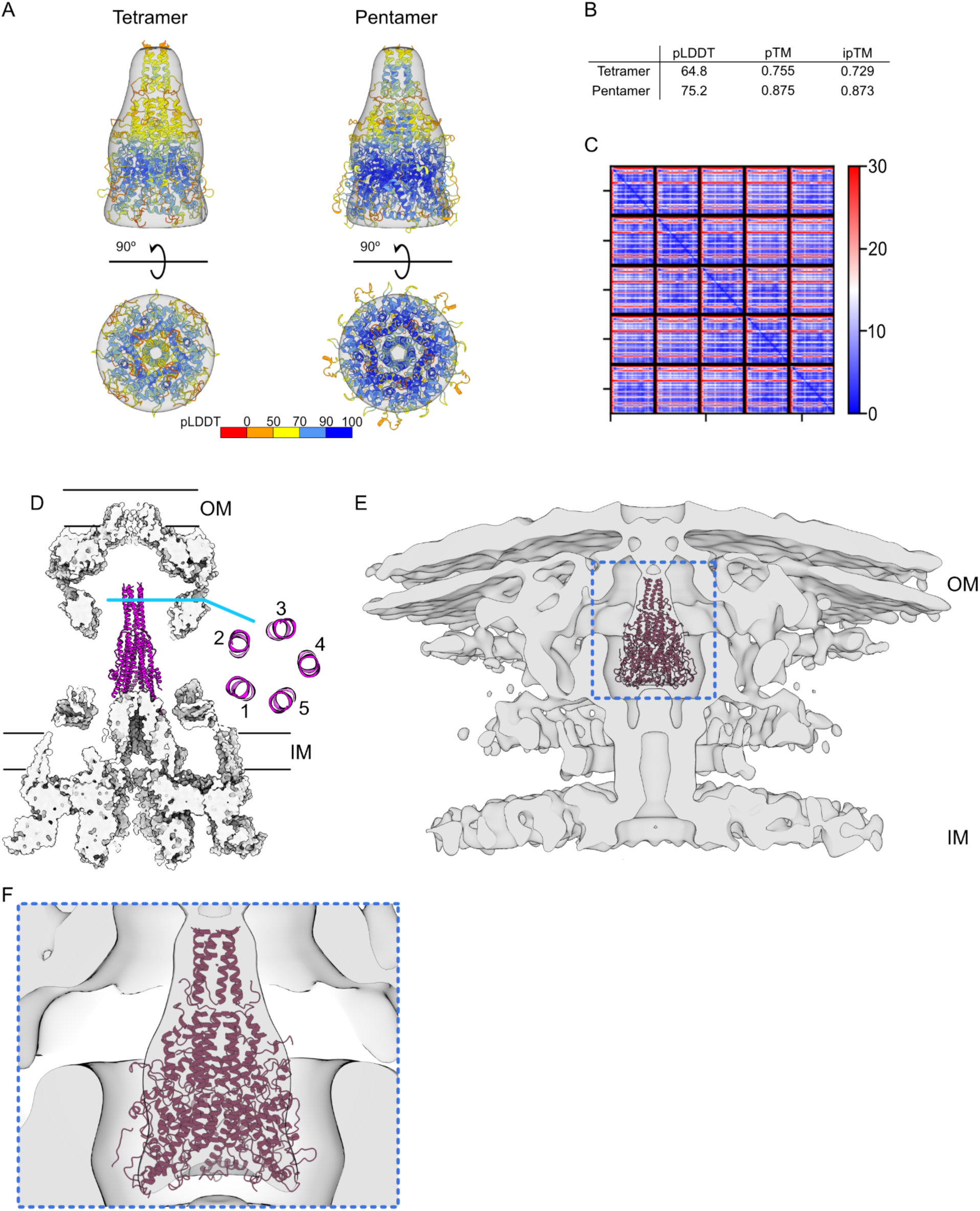
Modeling IcmX into the plug density. (**A**) AlphaFold/ColabFold predictions for the tetramer and pentamer of IcmX colored by pLDDT fit into the segmented plug density. (**B**) Modeling scores as returned by ColabFold for the highest-ranked models. pLDDT - predicted local distance difference test, representing local accuracy; pTM - predicted TM-score, corresponding to overall topological accuracy; ipTM - interface pTM score. (**C**) Heatmap represents the inter-chain accuracy of the model for the highest-rank pentameric model. Each pixel in the heatmap indicates alignment error in Angstroms for a pair of two residues. The metric indicates the positional uncertainty at residue x if the predicted and actual structures are aligned on residue y (*88*). Blue represents low error. (**D**) Atomic model of the conjugative T4SS (PBD 7O3J, 7O3T, 7O3V, 7O41) (*17*) with the VirB5 pentamer forming a stalk with structural similarity to IcmX. (**E**) Rigid-body fitting of the pentameric model of IcmX into the cryo-ET map. (**F**) Enlarged view as indicated by the blue dashed outline in panel **E**.

**Figure S11.**
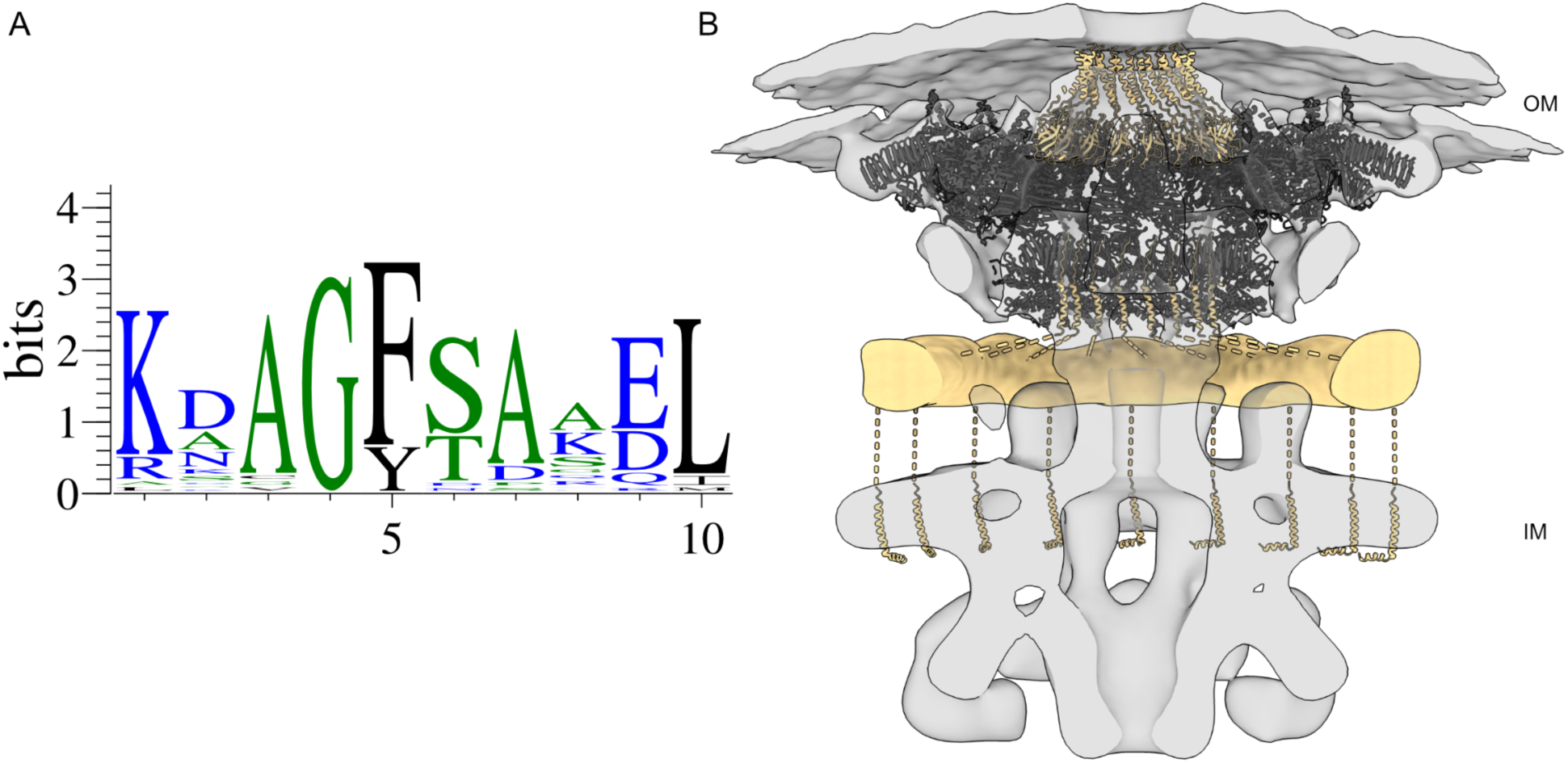
DotG as potential collar forming protein. (**A**) Weblogo plot showing consensus sequence of the 10 amino-acid repeating sequence in DotG. (**B**) Schematic representation showing how DotG could continue from the PR to form the collar and eventually anchor in the IM. Atomic model of the OMC and PR (PDB 7MUS) (*16*) in black. DotG highlighted in beige.

**Figure S12.**
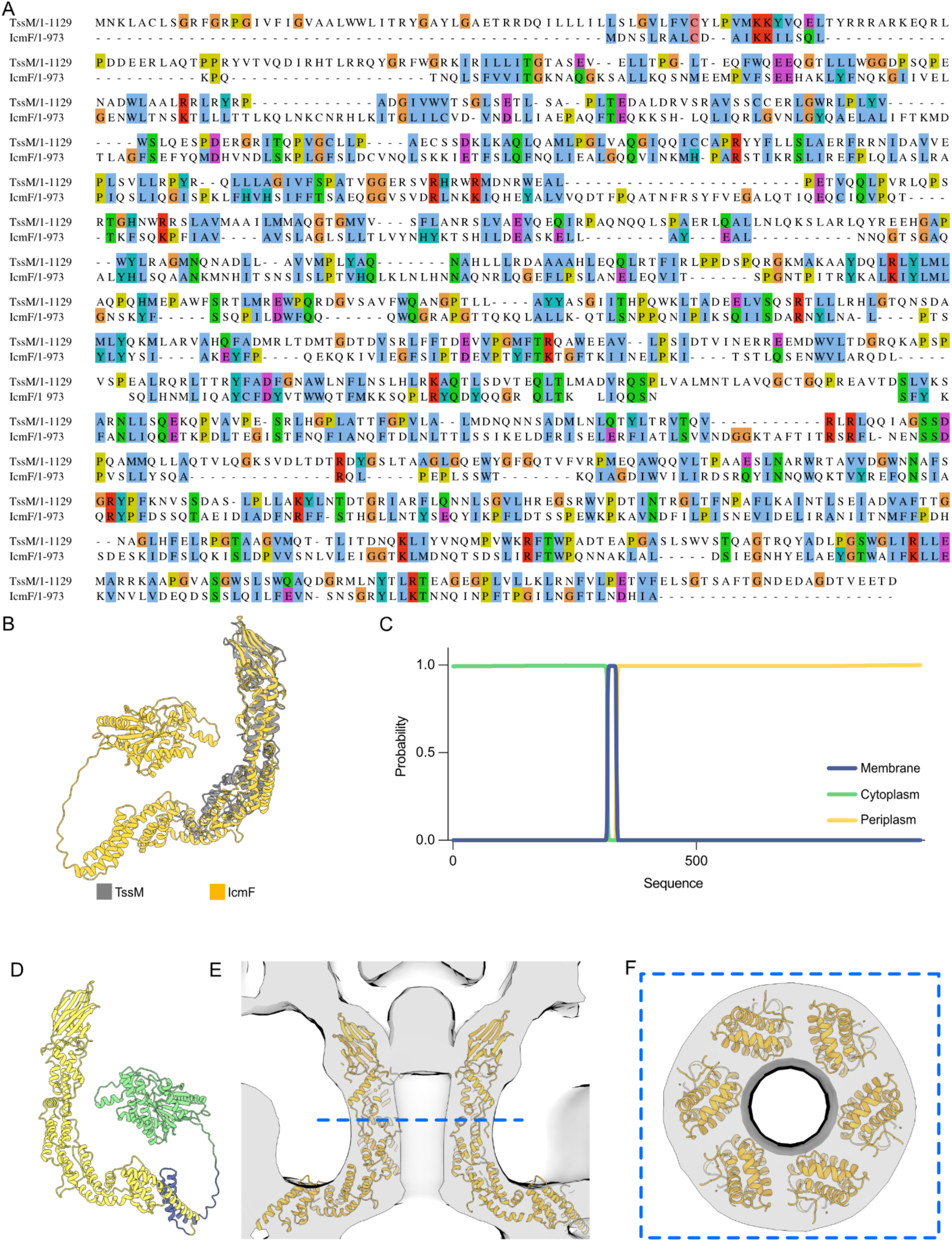
Modeling IcmF into the cylinder density. (**A**) Protein sequence alignment between TssM (T6SS) and IcmF. (**B**) Structure alignment between the structure of one of the TssM monomers that form the T6SS core complex channel (PDB 6IXH) (*34*) and a monomer of IcmF as predicted by AlphaFold. (**C**) IcmF topology prediction by DeepTMHMM (*26*). (**D**) IcmF model colored according to the predicted topology. (**E**-**F**) Rigid-body fitting of the IcmF hexamer into the cylinder density of the cryo-ET map. (**F**) Cross-section at the position indicated by the blue dashed line in panel **E**.

**Figure S13.**
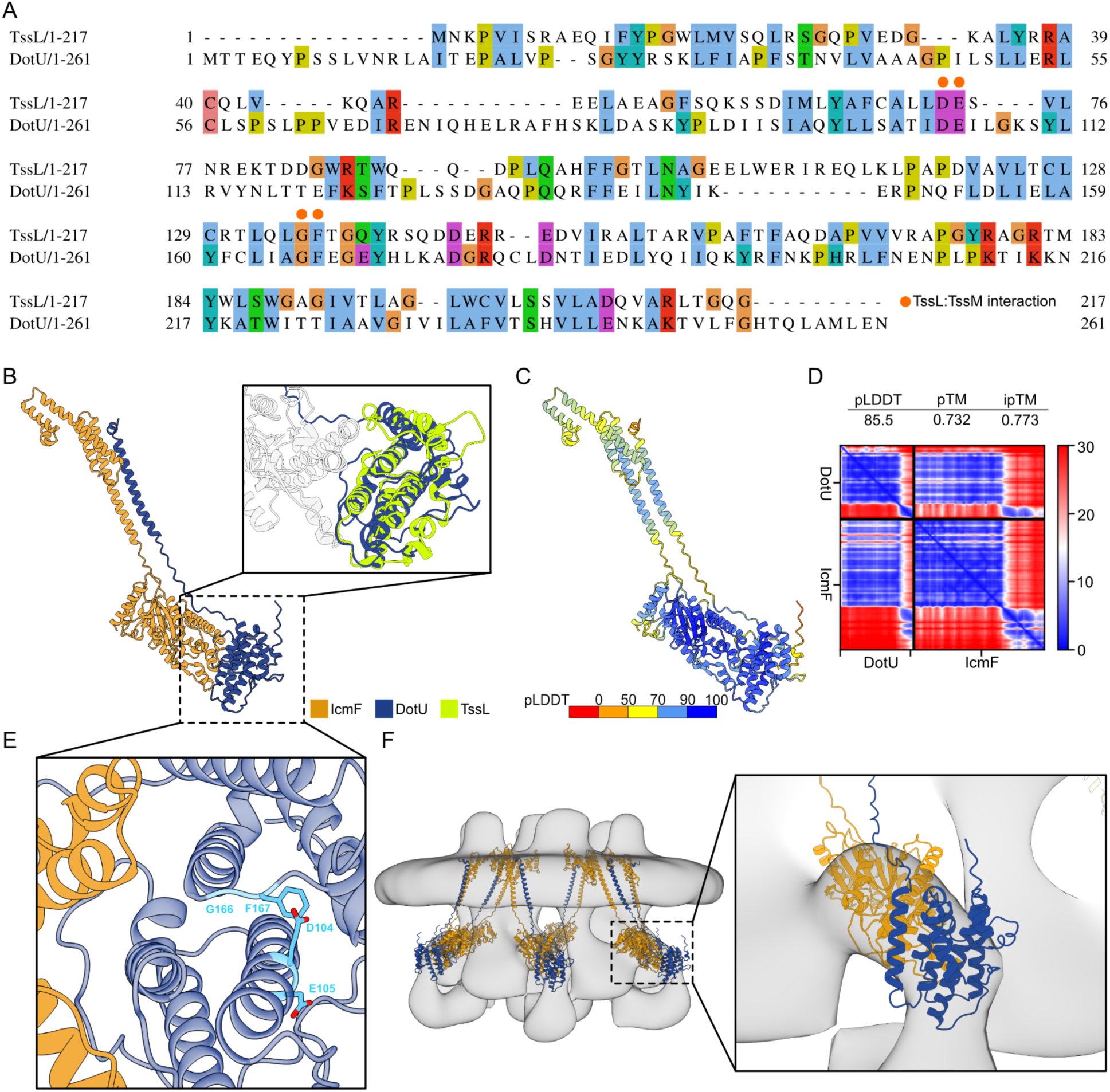
Modeling the IcmF_cyto_:DotU subcomplex. **(A)** Protein sequence alignment between TssL (T6SS) and DotU. Orange dots indicate the location of the residues important for TssM:TssL interaction in T6SS (*29*). These residues are conserved in DotU. (**B**) AlphaFold/ColabFold prediction of the IcmF_cyto_:DotU heterodimer. The inset shows the structural similarity between the DotU model and the crystal structure of TssL (PDB 3U66) (*36*). (**C**) IcmF_cyto_:DotU model colored by pLDDT. (**D**) Modeling scores as returned by ColabFold for the highest-ranked model. For a full description see Fig. S10. (**E**) Location of conserved residues in DotU identified previously as important for the TssM:TssL interaction (*29*). Residues are seen here to be located close to the IcmF_cyto_:DotU interaction interface. Our interpretation is that point mutations disrupt tertiary structure and therefore the IcmF_cyto_:DotU interaction. (**F**) Docking of the IcmF_cyto_:DotU subcomplexes into the top of the comma density in the cryo-ET map.

**Figure S14.**
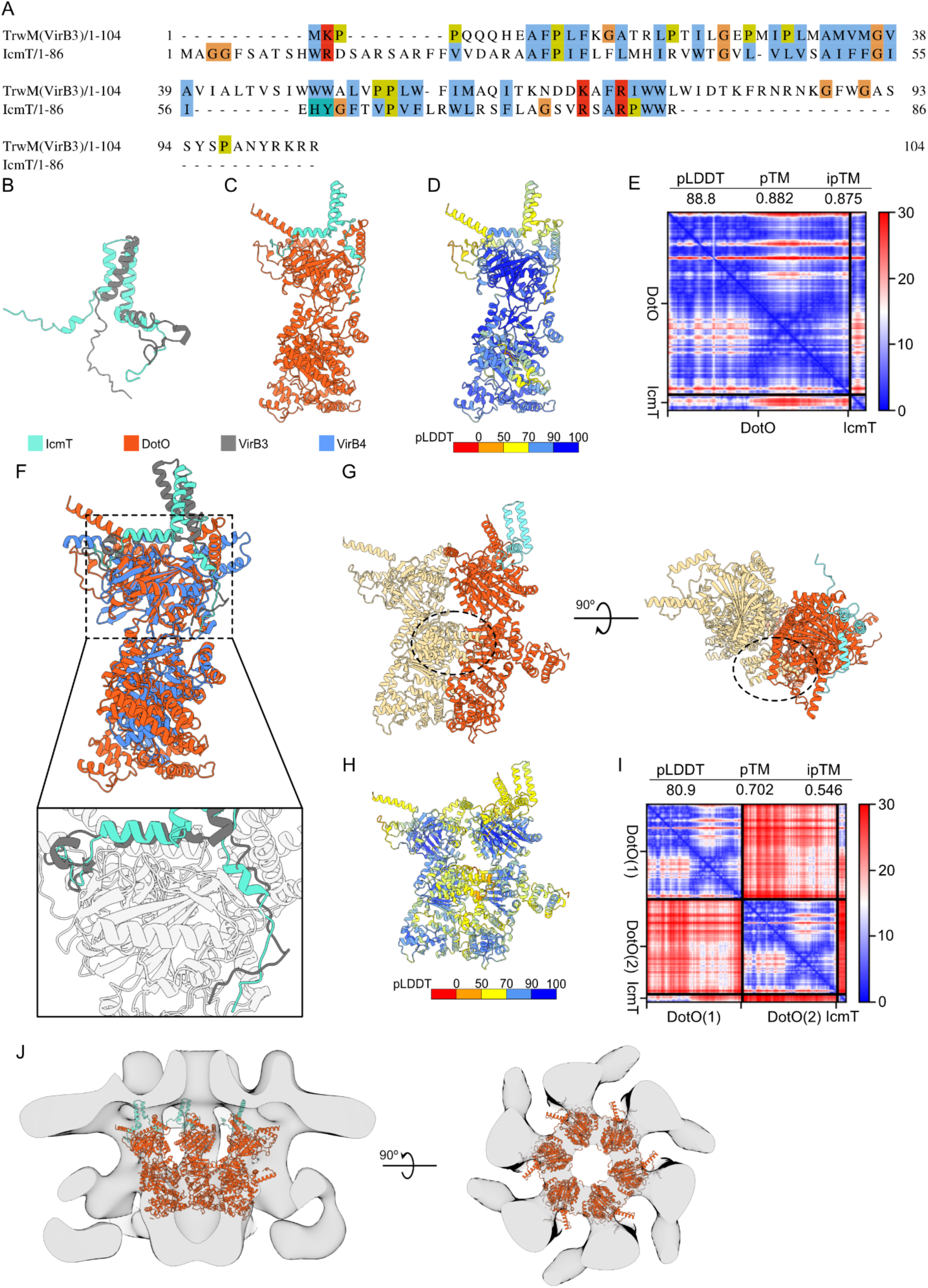
Modeling the IcmT:DotO ATPase subcomplexes. (**A**) Protein sequence alignment between VirB3 and IcmT. (**B**) Structure alignment between the experimentally determined structure of VirB3 (PDB 7O41) (*17*) and AlphaFold prediction for IcmT. (**C**-**D**) Predicted structure of IcmT:DotO subcomplex from AlphaFold/ColabFold. (**D**) IcmT:DotO model colored by pLDDT. (**E**) Modeling scores as returned by ColabFold for the highest-ranked model. For a full description see Fig. S10. (**F**) Overlay of the IcmT:DotO predicted model with the experimental structure of homologous VirB3/VirB4 (PDB 7O41) (*17*). The inset shows the conserved binding site of IcmT and VirB3. (**G-I**) AlphaFold/ColabFold model for DotO dimer with IcmT. (**G**) The C-terminal portion responsible for inter-subunit interaction between DotO is indicated by a dashed circle. (**H**) Predicted model colored by pLDDT. (**I**) Modeling scores as returned by ColabFold for the highest-ranked model. For a full description see Fig. S10. (**J**) IcmT:DotO inner hexamer fit into the cryo-ET map (gray).

**Figure S15.**
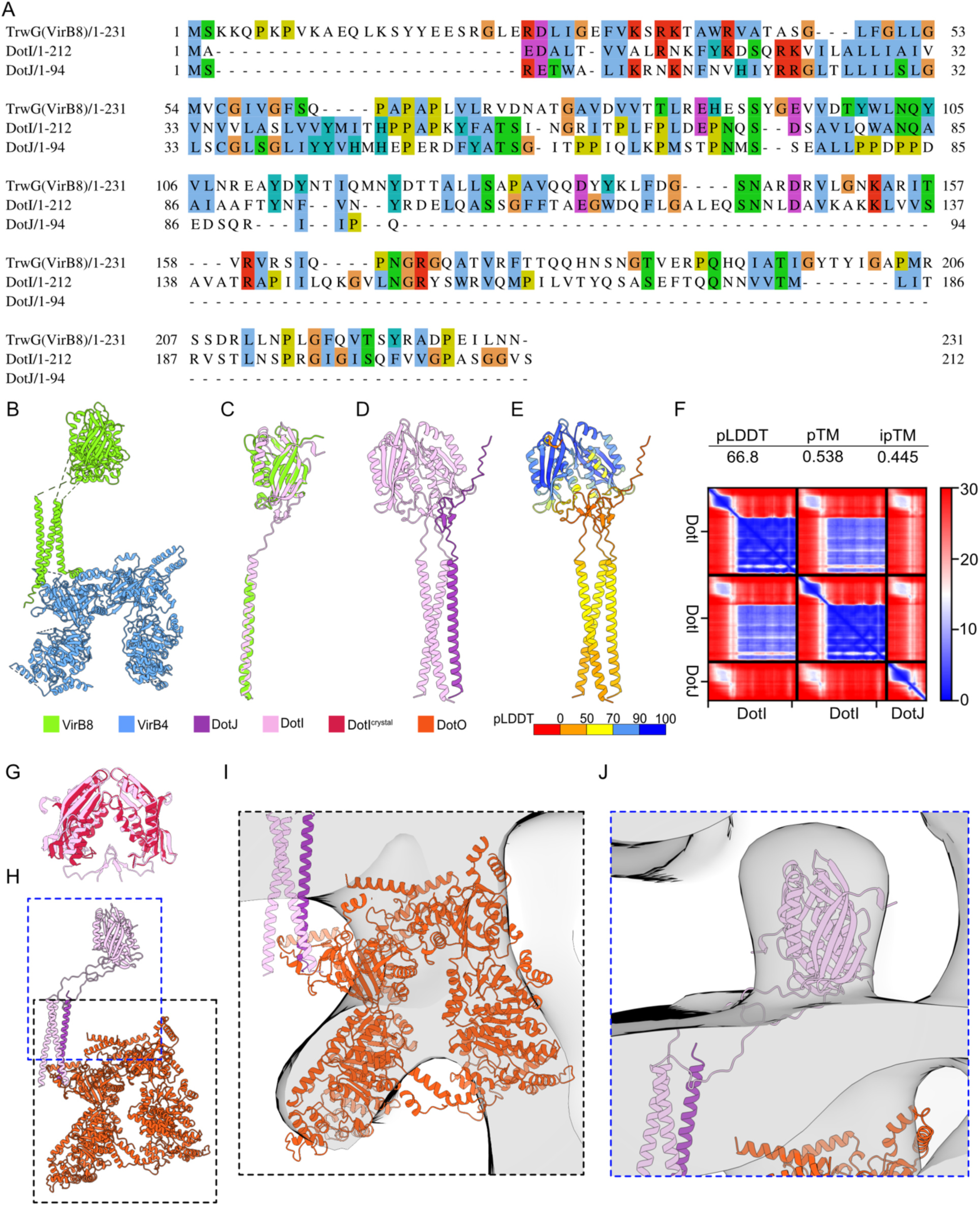
Modeling the DotI:DotJ:DotO subcomplex. (**A**) Protein sequence alignment between VirB8, DotI, and DotJ. (**B**) Structure of the VirB8:VirB4 subcomplex (PDB 7O41) (*17*). (**C**) Structure alignment between the experimentally determined structure of VirB8 (PDB 7O41) (*17*) and AlphaFold prediction for DotI. (**D**) 2:1 DotI:DotJ subcomplex structure predicted by ColabFold/AlphaFold. (**E**) DotI:DotJ model colored by pLDDT. (**F**) Modeling scores as returned by ColabFold for the highest-ranked model. (**G**) Alignment of the periplasmic portion of the DotI dimers comparing the AlphaFold prediction and a crystal structure (PDB 3WZ5) (*31*). (**H**) Subcomplex of DotO dimer with DotI:DotJ trimer modeled based on the VirB8:VirB4 structure. (**I**) The cytosolic portion of the DotI:DotJ:DotO subcomplex fit into the cryo-ET map. (**J**) DotI periplasmic portion fit into the cryo-ET map.

**Figure S16.**
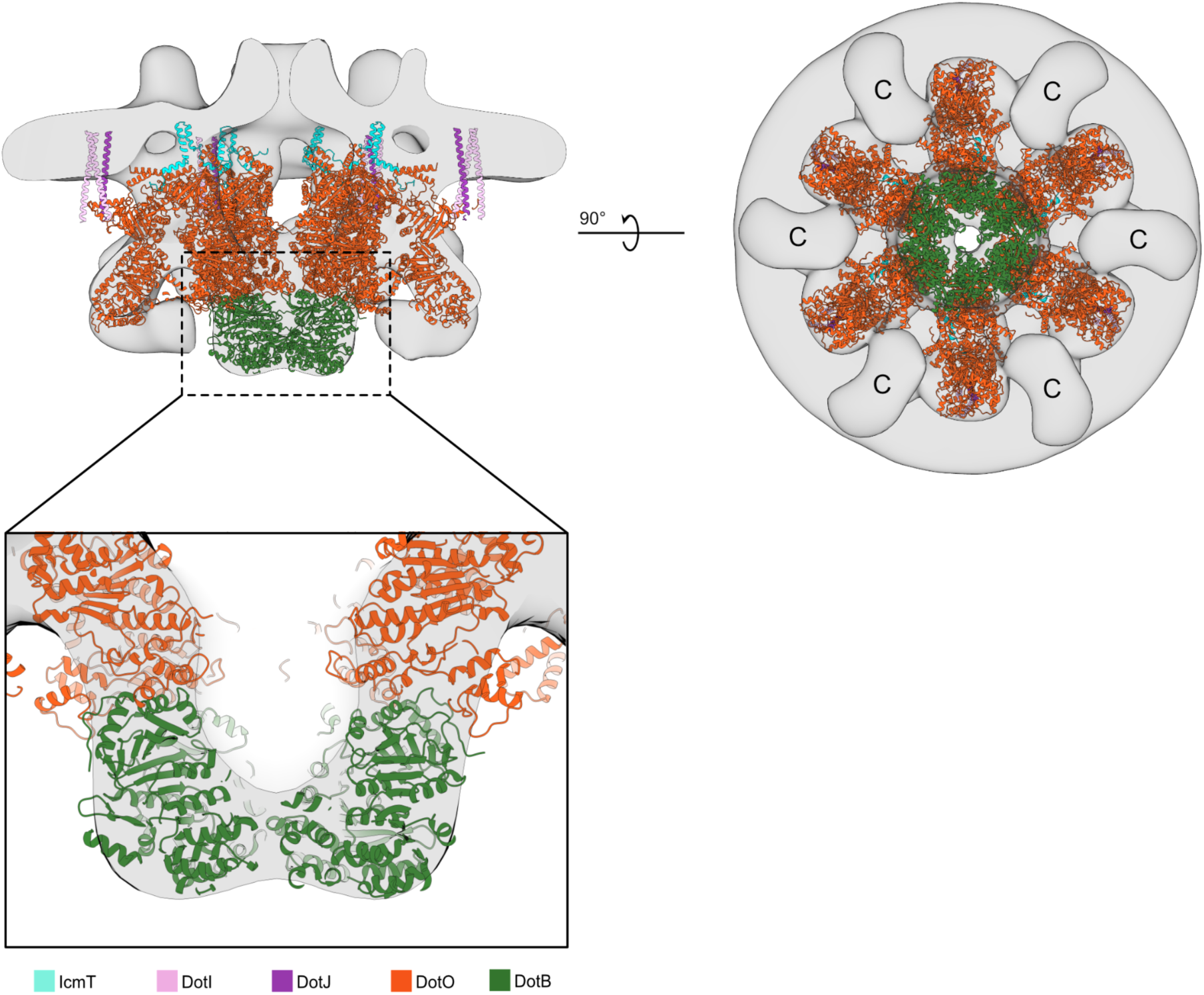
Placement of DotB. DotB crystal structure (PDB 6GEB) (*32*) docked into the density below the inner DotO hexamer in the cryo-ET map. The fully assembled ATPases (DotO:DotB) are anchored to the IM through IcmT (inner DotO) and DotI:DotJ (outer DotO). C - comma density.

**Figure S17.**
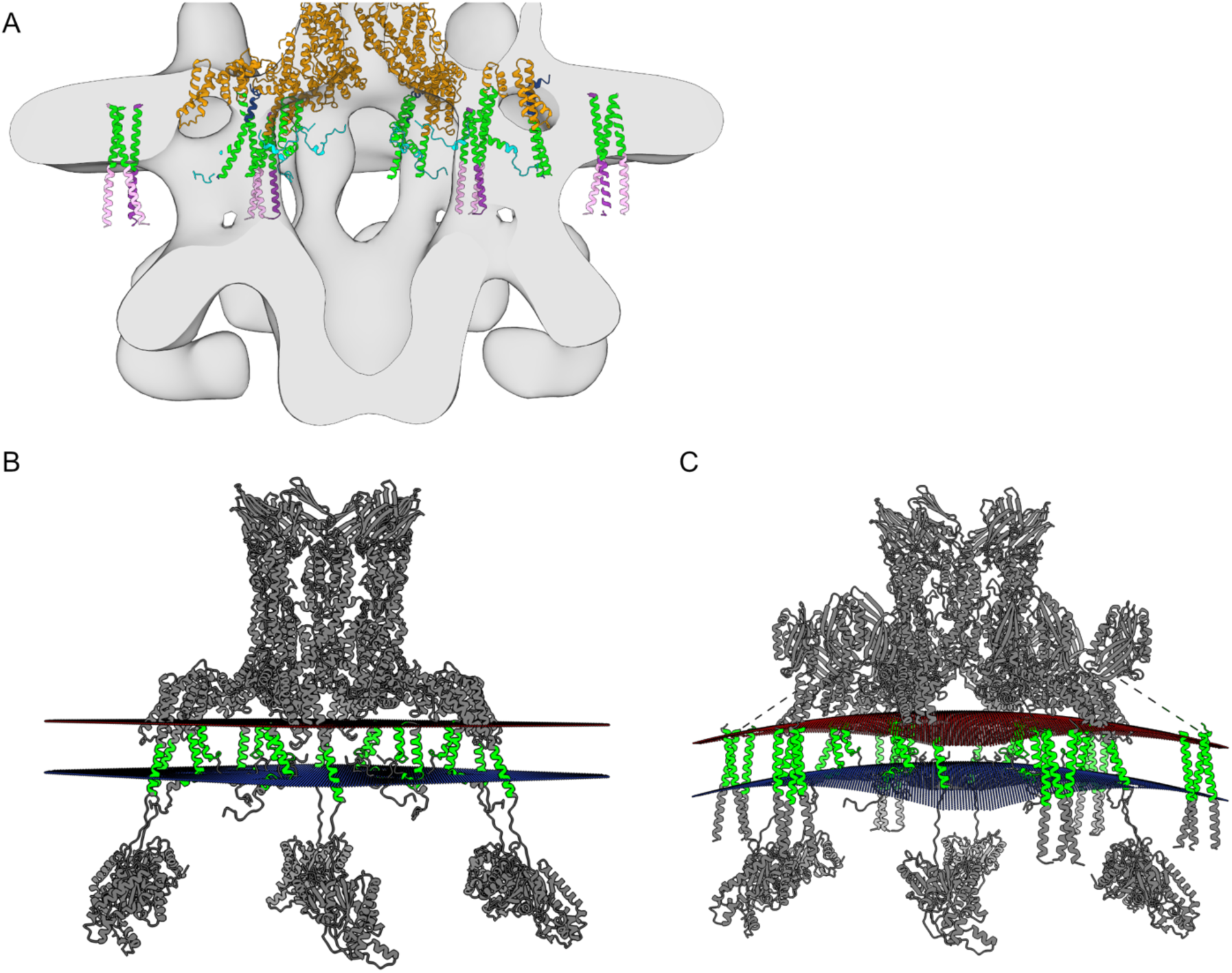
Localization of the transmembrane domains in the IM. (**A**) Model of the IMC fit into the cryo-ET map. Cytosolic portions of the proteins are not shown. (**B**,**C**) Simulated membrane topology using PP3 package for the model with IcmT:IcmF complex (**B**) and IcmF:IcmT:DotI:DotJ complex (**C**). Predicted transmembrane regions are highlighted in green.

**Figure S18.**
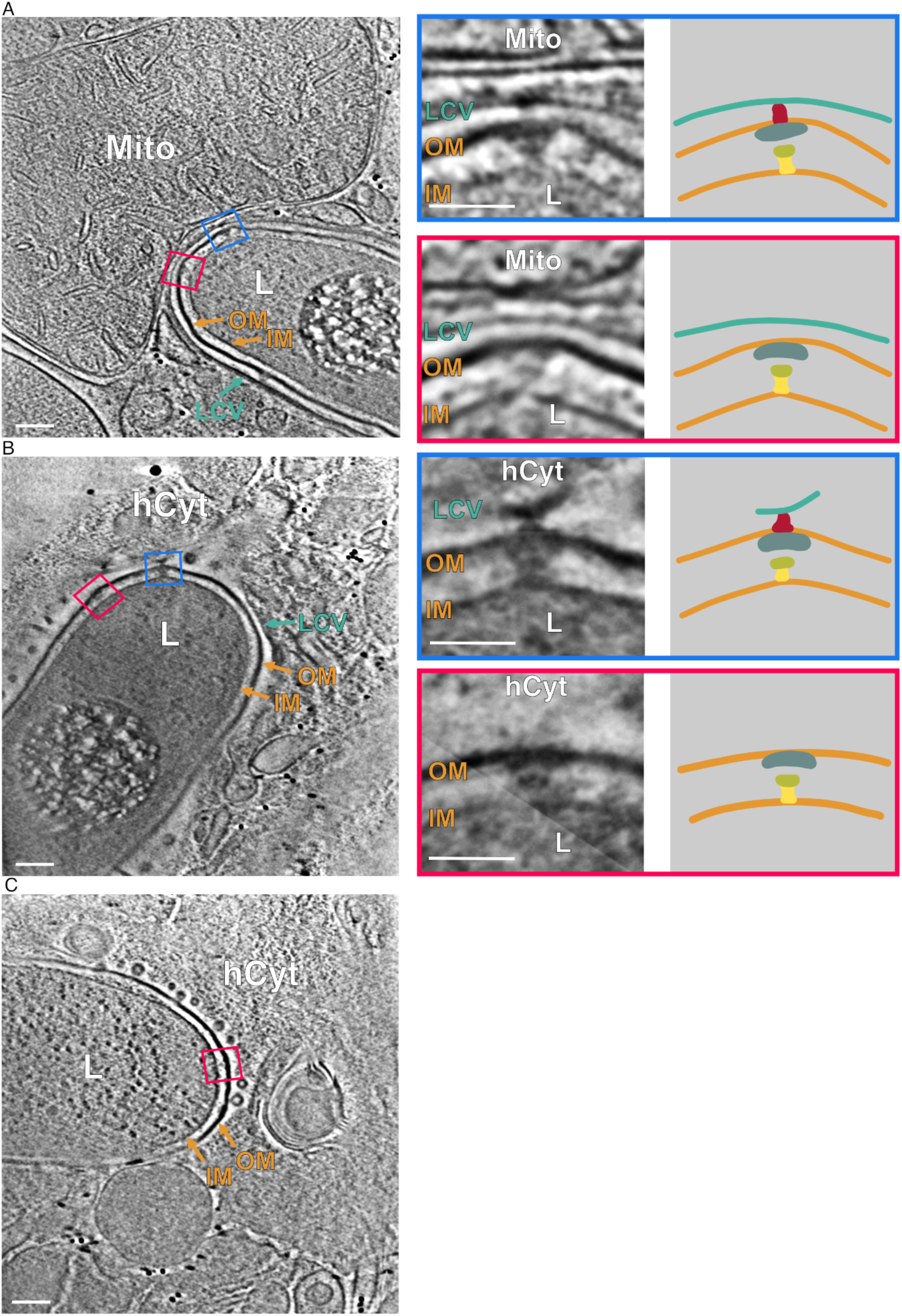
Additional examples of *L. pneumophila* inside host cells during early infection. (**A-C**) Representative tomographic slices of intracellular Legionella imaged at 1 h post-infection. OM (orange) - *L. pneumophila* outer membrane; IM (orange) - *L. pneumophila* inner membrane; LCV (cyan) - *L. pneumophila* containing vacuoles. L (orange) - *L. pneumophila* cells; hER (brown) - endoplasmic reticulum of U937 cells. Particles tethered to the host membrane are enclosed by blue frames; particles without a tethering site are indicated by red frames. The panels on the right depict individual particles making the OMC, PR, and cylinder schematically. Scale bars, 100 nm for main panels and 50 nm for the insets.

**Table S1.**
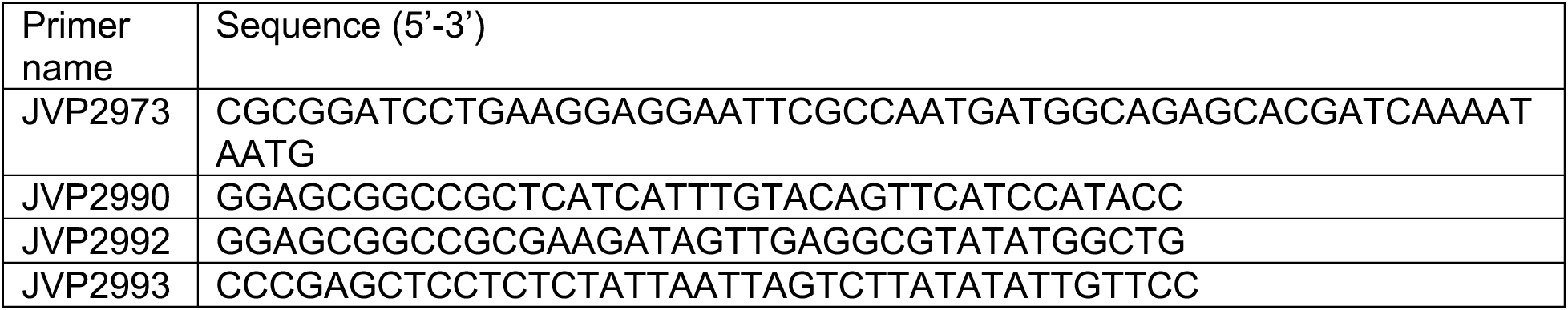
Primers used for L. pneumophila strains preparation.

## Notes

### Competing Interest Statement

The authors have declared no competing interest.

## REFERENCES

1. S. Backert, E. Grohmann, Type IV Secretion in Gram-Negative and Gram-Positive Bacteria (Springer, 2018; https://play.google.com/store/books/details?id=CjtRDwAAQBAJ).

2. Y. G. Li, B. Hu, P. J. Christie, Biological and Structural Diversity of Type IV Secretion Systems. Microbiol Spectr. 7 (2019), doi:10.1128/microbiolspec.PSIB-0012-2018.

3. D. Dubnau, M. Blokesch, Mechanisms of DNA Uptake by Naturally Competent Bacteria. Annu. Rev. Genet. 53, 217–237 (2019).

4. K. L. Dooling, K.-A. Toews, L. A. Hicks, L. E. Garrison, B. Bachaus, S. Zansky, L. R. Carpenter, B. Schaffner, E. Parker, S. Petit, A. Thomas, S. Thomas, R. Mansmann, C. Morin, B. White, G. E. Langley, Active Bacterial Core Surveillance for Legionellosis - United States, 2011-2013. MMWR Morb. Mortal. Wkly. Rep. 64, 1190–1193 (2015).

5. A. Hubber, C. R. Roy, Modulation of host cell function by Legionella pneumophila type IV effectors. Annu. Rev. Cell Dev. Biol. 26, 261–283 (2010).

6. J. Qiu, Z.-Q. Luo, Legionella and Coxiella effectors: strength in diversity and activity. Nat. Rev. Microbiol. 15, 591–605 (2017).

7. T. Kubori, H. Nagai, The Type IVB secretion system: an enigmatic chimera. Curr. Opin. Microbiol. 29, 22–29 (2016).

8. D. Ghosal, Y. Chang, K. C. Jeong, J. P. Vogel, G. J. Jensen, *In situ* structure of the *Legionella* Dot/Icm type IV secretion system by electron cryotomography. EMBO reports. 18 (2017), pp. 726–732.

9. Y.-W. Chang, C. L. Shaffer, L. A. Rettberg, D. Ghosal, G. J. Jensen, In Vivo Structures of the Helicobacter pylori cag Type IV Secretion System. Cell Rep. 23, 673–681 (2018).

10. D. Ghosal, K. C. Jeong, Y.-W. Chang, J. Gyore, L. Teng, A. Gardner, J. P. Vogel, G. J. Jensen, Molecular architecture, polar targeting and biogenesis of the Legionella Dot/Icm T4SS. Nature Microbiology. 4 (2019), pp. 1173–1182.

11. M. Kaplan, D. C. Shepherd, N. Vankadari, K. W. Kim, C. L. Larson, P. Dutka, P. A. Beare, E. Krzymowski, R. A. Heinzen, G. J. Jensen, D. Ghosal, Structural remodeling of Coxiella burnetii during its biphasic developmental cycle revealed by cryo-electron tomography. bioRxiv (2022), p. 2022.08.23.505044.

12. D. Chetrit, B. Hu, P. J. Christie, C. R. Roy, J. Liu, A unique cytoplasmic ATPase complex defines the Legionella pneumophila type IV secretion channel. Nature Microbiology. 3 (2018), pp. 678–686.

13. D. Park, D. Chetrit, B. Hu, C. R. Roy, J. Liu, Analysis of Dot/Icm Type IVB Secretion System Subassemblies by Cryoelectron Tomography Reveals Conformational Changes Induced by DotB Binding. MBio. 11 (2020), doi:10.1128/mBio.03328-19.

14. J. M. Chung, M. J. Sheedlo, A. M. Campbell, N. Sawhney, A. E. Frick-Cheng, D. B. Lacy, T. L. Cover, M. D. Ohi, Structure of the Helicobacter pylori Cag type IV secretion system. Elife. 8 (2019), doi:10.7554/eLife.47644.

15. C. L. Durie, M. J. Sheedlo, J. M. Chung, B. G. Byrne, M. Su, T. Knight, M. Swanson, D. B. Lacy, M. D. Ohi, Structural analysis of the Legionella pneumophila Dot/Icm type IV secretion system core complex. Elife. 9 (2020), doi:10.7554/eLife.59530.

16. M. J. Sheedlo, C. L. Durie, J. M. Chung, L. Chang, J. Roberts, M. Swanson, D. B. Lacy, M. D. Ohi, Cryo-EM reveals new species-specific proteins and symmetry elements in the Legionella pneumophila Dot/Icm T4SS. eLife. 10 (2021),, doi:10.7554/elife.70427.

17. K. Macé, A. K. Vadakkepat, A. Redzej, N. Lukoyanova, C. Oomen, N. Braun, M. Ukleja, F. Lu, T. R. D. Costa, E. V. Orlova, D. Baker, Q. Cong, G. Waksman, Cryo-EM structure of a type IV secretion system. Nature. 607, 191–196 (2022).

18. S. M. VanRheenen, G. Duménil, R. R. Isberg, IcmF and DotU are required for optimal effector translocation and trafficking of the Legionella pneumophila vacuole. Infect. Immun. 72, 5972–5982 (2004).

19. J. A. Sexton, H.-J. Yeo, J. P. Vogel, Genetic analysis of the Legionella pneumophila DotB ATPase reveals a role in type IV secretion system protein export. Mol. Microbiol. 57, 70–84 (2005).

20. H. Winkler, 3D reconstruction and processing of volumetric data in cryo-electron tomography. J. Struct. Biol. 157, 126–137 (2007).

21. J. Jumper, R. Evans, A. Pritzel, T. Green, M. Figurnov, O. Ronneberger, K. Tunyasuvunakool, R. Bates, A. Žídek, A. Potapenko, A. Bridgland, C. Meyer, S. A. A. Kohl, A. J. Ballard, A. Cowie, B. Romera-Paredes, S. Nikolov, R. Jain, J. Adler, T. Back, S. Petersen, D. Reiman, E. Clancy, M. Zielinski, M. Steinegger, M. Pacholska, T. Berghammer, S. Bodenstein, D. Silver, O. Vinyals, A. W. Senior, K. Kavukcuoglu, P. Kohli, D. Hassabis, Highly accurate protein structure prediction with AlphaFold. Nature. 596, 583– 589 (2021).

22. C. D. Vincent, J. R. Friedman, K. C. Jeong, E. C. Buford, J. L. Miller, J. P. Vogel, Identification of the core transmembrane complex of the Legionella Dot/Icm type IV secretion system. Mol. Microbiol. 62, 1278–1291 (2006).

23. M. Mirdita, K. Schütze, Y. Moriwaki, L. Heo, S. Ovchinnikov, M. Steinegger, ColabFold: making protein folding accessible to all. Nat. Methods (2022), doi:10.1038/s41592-022-01488-1.

24. L. Gomez-Valero, A. Chiner-Oms, I. Comas, C. Buchrieser, Evolutionary Dissection of the Dot/Icm System Based on Comparative Genomics of 58 Legionella Species. Genome Biol. Evol. 11, 2619–2632 (2019).

25. H. Nagai, C. R. Roy, The DotA protein from Legionella pneumophila is secreted by a novel process that requires the Dot/Icm transporter. EMBO J. 20, 5962–5970 (2001).

26. J. Hallgren, K. D. Tsirigos, M. D. Pedersen, J. J. A. Armenteros, P. Marcatili, H. Nielsen, A. Krogh, O. Winther, DeepTMHMM predicts alpha and beta transmembrane proteins using deep neural networks. bioRxiv (2022), p. 2022.04.08.487609.

27. L.-S. Ma, F. Narberhaus, E.-M. Lai, IcmF family protein TssM exhibits ATPase activity and energizes type VI secretion. J. Biol. Chem. 287, 15610–15621 (2012).

28. L. Logger, M.-S. Aschtgen, M. Guérin, E. Cascales, E. Durand, Molecular Dissection of the Interface between the Type VI Secretion TssM Cytoplasmic Domain and the TssG Baseplate Component. J. Mol. Biol. 428, 4424–4437 (2016).

29. A. Zoued, C. J. Cassaro, E. Durand, B. Douzi, A. P. España, C. Cambillau, L. Journet, E. Cascales, Structure-Function Analysis of the TssL Cytoplasmic Domain Reveals a New Interaction between the Type VI Secretion Baseplate and Membrane Complexes. J. Mol. Biol. 428, 4413–4423 (2016).

30. J. A. Sexton, J. L. Miller, A. Yoneda, T. E. Kehl-Fie, J. P. Vogel, Legionella pneumophila DotU and IcmF are required for stability of the Dot/Icm complex. Infect. Immun. 72, 5983– 5992 (2004).

31. T. Kuroda, T. Kubori, X. Thanh Bui, A. Hyakutake, Y. Uchida, K. Imada, H. Nagai, Molecular and structural analysis of Legionella DotI gives insights into an inner membrane complex essential for type IV secretion. Sci. Rep. 5, 10912 (2015).

32. M. S. Prevost, G. Waksman, X-ray crystal structures of the type IVb secretion system DotB ATPases. Protein Science. 27 (2018), pp. 1464–1475.

33. C. Rapisarda, Y. Cherrak, R. Kooger, V. Schmidt, R. Pellarin, L. Logger, E. Cascales, M. Pilhofer, E. Durand, R. Fronzes, In situ and high-resolution cryo-EM structure of a bacterial type VI secretion system membrane complex. EMBO J. 38 (2019), doi:10.15252/embj.2018100886.

34. M. Yin, Z. Yan, X. Li, Architecture of type VI secretion system membrane core complex. Cell Res. 29, 251–253 (2019).

35. L. Renault, B. Guibert, J. Cherfils, Structural snapshots of the mechanism and inhibition of a guanine nucleotide exchange factor. Nature. 426, 525–530 (2003).

36. E. Durand, A. Zoued, S. Spinelli, P. J. H. Watson, M.-S. Aschtgen, L. Journet, C. Cambillau, E. Cascales, Structural characterization and oligomerization of the TssL protein, a component shared by bacterial type VI and type IVb secretion systems. J. Biol. Chem. 287, 14157–14168 (2012).

37. P. Escoll, O.-R. Song, F. Viana, B. Steiner, T. Lagache, J.-C. Olivo-Marin, F. Impens, P. Brodin, H. Hilbi, C. Buchrieser, Legionella pneumophila Modulates Mitochondrial Dynamics to Trigger Metabolic Repurposing of Infected Macrophages. Cell Host Microbe. 22, 302– 316.e7 (2017).

38. D. Böck, D. Hüsler, B. Steiner, J. M. Medeiros, A. Welin, K. A. Radomska, W.-D. Hardt, M. Pilhofer, H. Hilbi, The Polar Legionella Icm/Dot T4SS Establishes Distinct Contact Sites with the Pathogen Vacuole Membrane. MBio. 12, e0218021 (2021).

39. A. Meir, K. Macé, N. Lukoyanova, D. Chetrit, M. K. Hospenthal, A. Redzej, C. Roy, G. Waksman, Mechanism of effector capture and delivery by the type IV secretion system from Legionella pneumophila. Nat. Commun. 11, 2864 (2020).

40. D. C. Lockwood, H. Amin, T. R. D. Costa, G. N. Schroeder, The Legionella pneumophila Dot/Icm type IV secretion system and its effectors. Microbiology. 168 (2022), doi:10.1099/mic.0.001187.

41. M. J. Sheedlo, M. D. Ohi, D. B. Lacy, T. L. Cover, Molecular architecture of bacterial type IV secretion systems. PLoS Pathog. 18, e1010720 (2022).

42. P. J. Christie, L. Gomez Valero, C. Buchrieser, Biological Diversity and Evolution of Type IV Secretion Systems. Curr. Top. Microbiol. Immunol. 413, 1–30 (2017).

43. V. S. Nguyen, B. Douzi, E. Durand, A. Roussel, E. Cascales, C. Cambillau, Towards a complete structural deciphering of Type VI secretion system. Curr. Opin. Struct. Biol. 49, 77–84 (2018).

44. J. Wang, M. Brodmann, M. Basler, Assembly and Subcellular Localization of Bacterial Type VI Secretion Systems. Annu. Rev. Microbiol. 73, 621–638 (2019).

45. Alvarez-Martinez Cristina E., Christie Peter J., Biological Diversity of Prokaryotic Type IV Secretion Systems. Microbiol. Mol. Biol. Rev. 73, 775–808 (2009).

46. E. Cascales, P. J. Christie, The versatile bacterial type IV secretion systems. Nat. Rev. Microbiol. 1, 137–149 (2003).

47. P. J. Christie, The Mosaic Type IV Secretion Systems. EcoSal Plus. 7 (2016), doi:10.1128/ecosalplus.ESP-0020-2015.

48. M. J. Sheedlo, J. M. Chung, N. Sawhney, C. L. Durie, T. L. Cover, M. D. Ohi, D. B. Lacy, Cryo-EM reveals species-specific components within the Helicobacter pylori Cag type IV secretion system core complex. Elife. 9 (2020), doi:10.7554/eLife.59495.

49. P. Khara, L. Song, P. J. Christie, B. Hu, In Situ Visualization of the pKM101-Encoded Type IV Secretion System Reveals a Highly Symmetric ATPase Energy Center. MBio. 12, e0246521 (2021).

50. B. Hu, P. Khara, P. J. Christie, Structural bases for F plasmid conjugation and F pilus biogenesis in *Escherichia coli*. Proc. Natl. Acad. Sci. U. S. A. 116, 14222–14227 (2019).

51. M. Matthews, C. R. Roy, Identification and subcellular localization of the Legionella pneumophila IcmX protein: a factor essential for establishment of a replicative organelle in eukaryotic host cells. Infect. Immun. 68, 3971–3982 (2000).

52. G. Segal, M. Feldman, T. Zusman, The Icm/Dot type-IV secretion systems of Legionella pneumophila and Coxiella burnetii. FEMS Microbiol. Rev. 29, 65–81 (2005).

53. W. M. Amyot, D. deJesus, R. R. Isberg, Poison domains block transit of translocated substrates via the Legionella pneumophila Icm/Dot system. Infect. Immun. 81, 3239–3252 (2013).

54. G. Chreifi, S. Chen, G. J. Jensen, Rapid tilt-series method for cryo-electron tomography: Characterizing stage behavior during FISE acquisition. J. Struct. Biol. 213, 107716 (2021).

55. G. Chreifi, S. Chen, L. A. Metskas, M. Kaplan, G. J. Jensen, Rapid tilt-series acquisition for electron cryotomography. J. Struct. Biol. 205, 163–169 (2019).

56. D. N. Mastronarde, Automated electron microscope tomography using robust prediction of specimen movements. J. Struct. Biol. 152, 36–51 (2005).

57. C. M. Oikonomou, G. J. Jensen, Cellular Electron Cryotomography: Toward Structural Biology In Situ. Annu. Rev. Biochem. 86, 873–896 (2017).

58. R. Evans, M. O’Neill, A. Pritzel, N. Antropova, A. Senior, T. Green, A. Žídek, R. Bates, S. Blackwell, J. Yim, O. Ronneberger, S. Bodenstein, M. Zielinski, A. Bridgland, A. Potapenko, A. Cowie, K. Tunyasuvunakool, R. Jain, E. Clancy, P. Kohli, J. Jumper, D. Hassabis, Protein complex prediction with AlphaFold-Multimer. bioRxiv (2022), p. 2021.10.04.463034.

59. W. F. Tivol, A. Briegel, G. J. Jensen, An improved cryogen for plunge freezing. Microsc. Microanal. 14, 375–379 (2008).

60. S. Q. Zheng, E. Palovcak, J.-P. Armache, K. A. Verba, Y. Cheng, D. A. Agard, MotionCor2: anisotropic correction of beam-induced motion for improved cryo-electron microscopy. Nat. Methods. 14, 331–332 (2017).

61. J. R. Kremer, D. N. Mastronarde, J. R. McIntosh, Computer visualization of three-dimensional image data using IMOD. J. Struct. Biol. 116, 71–76 (1996).

62. M. Chen, J. M. Bell, X. Shi, S. Y. Sun, Z. Wang, S. J. Ludtke, A complete data processing workflow for cryo-ET and subtomogram averaging. Nat. Methods. 16, 1161–1168 (2019).

63. D. Castaño-Díez, M. Kudryashev, M. Arheit, H. Stahlberg, Dynamo: a flexible, user-friendly development tool for subtomogram averaging of cryo-EM data in high-performance computing environments. J. Struct. Biol. 178, 139–151 (2012).

64. J.-I. Agulleiro, J.-J. Fernandez, Tomo3D 2.0--exploitation of advanced vector extensions (AVX) for 3D reconstruction. J. Struct. Biol. 189, 147–152 (2015).

65. A. Kucukelbir, F. J. Sigworth, H. D. Tagare, Quantifying the local resolution of cryo-EM density maps. Nat. Methods. 11, 63–65 (2014).

66. T. D. Goddard, C. C. Huang, E. C. Meng, E. F. Pettersen, G. S. Couch, J. H. Morris, T. E. Ferrin, UCSF ChimeraX: Meeting modern challenges in visualization and analysis. Protein Sci. 27, 14–25 (2018).

67. V. Lam, E. Villa, Practical Approaches for Cryo-FIB Milling and Applications for Cellular Cryo-Electron Tomography. Methods Mol. Biol. 2215, 49–82 (2021).

68. D. N. Mastronarde, S. R. Held, Automated tilt series alignment and tomographic reconstruction in IMOD. J. Struct. Biol. 197, 102–113 (2017).

69. Y.-T. Liu, H. Zhang, H. Wang, C.-L. Tao, G.-Q. Bi, Z. H. Zhou, Isotropic reconstruction for electron tomography with deep learning. Nat. Commun. 13, 6482 (2022).

70. G. D. Pintilie, J. Zhang, T. D. Goddard, W. Chiu, D. C. Gossard, Quantitative analysis of cryo-EM density map segmentation by watershed and scale-space filtering, and fitting of structures by alignment to regions. J. Struct. Biol. 170, 427–438 (2010).

71. G. Pintilie, W. Chiu, Comparison of Segger and other methods for segmentation and rigid-body docking of molecular components in cryo-EM density maps. Biopolymers. 97, 742– 760 (2012).

72. A. Sali, T. L. Blundell, Comparative protein modelling by satisfaction of spatial restraints. J. Mol. Biol. 234, 779–815 (1993).

73. O. V. Stepanenko, O. V. Stepanenko, I. M. Kuznetsova, V. N. Uversky, K. K. Turoverov, Peculiarities of the Super-Folder GFP Folding in a Crowded Milieu. Int. J. Mol. Sci. 17 (2016), doi:10.3390/ijms17111805.

74. J. Y. Choi, T.-H. Jang, H. H. Park, The mechanism of folding robustness revealed by the crystal structure of extra-superfolder GFP. FEBS Lett. 591, 442–447 (2017).

75. M. Kanehisa, Linking databases and organisms: GenomeNet resources in Japan. Trends Biochem. Sci. 22, 442–444 (1997).

76. R. C. Edgar, MUSCLE: multiple sequence alignment with high accuracy and high throughput. Nucleic Acids Res. 32, 1792–1797 (2004).

77. R. C. Edgar, MUSCLE: a multiple sequence alignment method with reduced time and space complexity. BMC Bioinformatics. 5, 113 (2004).

78. G. E. Crooks, G. Hon, J.-M. Chandonia, S. E. Brenner, WebLogo: a sequence logo generator. Genome Res. 14, 1188–1190 (2004).

79. C. S. Robb, F. E. Nano, A. B. Boraston, The structure of the conserved type six secretion protein TssL (DotU) from Francisella novicida. J. Mol. Biol. 419, 277–283 (2012).

80. F. M. Ruiz, J. Lopez, C. G. Ferrara, E. Santillana, Y. R. Espinosa, M. F. Feldman, A. Romero, Structural Characterization of TssL from Acinetobacter baumannii: a Key Component of the Type VI Secretion System. J. Bacteriol. 202 (2020), doi:10.1128/JB.00210-20.

81. J. H. Chang, Y.-G. Kim, Crystal structure of the bacterial type VI secretion system component TssL from Vibrio cholerae. J. Microbiol. 53, 32–37 (2015).

82. G. C.P.van Zundert, Bijvoet Center for Biomolecular Research, Faculty of Science-Chemistry, Utrecht University, Utrecht, the Netherlands, A. M.J.J. Bonvin, Fast and sensitive rigid-body fitting into cryo-EM density maps with PowerFit. AIMS Biophys. 2, 73– 87 (2015).

83. A. L. Lomize, S. C. Todd, I. D. Pogozheva, Spatial arrangement of proteins in planar and curved membranes by PPM 3.0. Protein Sci. 31, 209–220 (2022).

84. F. Sievers, A. Wilm, D. Dineen, T. J. Gibson, K. Karplus, W. Li, R. Lopez, H. McWilliam, M. Remmert, J. Söding, Others, Fast, scalable generation of high-quality protein multiple sequence alignments using Clustal Omega. Mol. Syst. Biol. 7, 539 (2011).

85. A. M. Waterhouse, J. B. Procter, D. M. A. Martin, M. Clamp, G. J. Barton, Jalview Version 2—a multiple sequence alignment editor and analysis workbench. Bioinformatics. 25, 1189–1191 (2009).

86. S. Hare, R. Bayliss, C. Baron, G. Waksman, A large domain swap in the VirB11 ATPase of Brucella suis leaves the hexameric assembly intact. J. Mol. Biol. 360, 56–66 (2006).

87. Y.-W. Chang, L. A. Rettberg, D. R. Ortega, G. J. Jensen, In vivo structures of an intact type VI secretion system revealed by electron cryotomography. EMBO Rep. 18, 1090–1099 (2017).

88. M. Varadi, S. Anyango, M. Deshpande, S. Nair, C. Natassia, G. Yordanova, D. Yuan, O. Stroe, G. Wood, A. Laydon, A. Žídek, T. Green, K. Tunyasuvunakool, S. Petersen, J. Jumper, E. Clancy, R. Green, A. Vora, M. Lutfi, M. Figurnov, A. Cowie, N. Hobbs, P. Kohli, G. Kleywegt, E. Birney, D. Hassabis, S. Velankar, AlphaFold Protein Structure Database: massively expanding the structural coverage of protein-sequence space with high-accuracy models. Nucleic Acids Res. 50, D439–D444 (2022).

